# Pre-clinical validation of an RNA-based precision oncology platform for patient-therapy alignment in a diverse set of human malignancies resistant to standard treatments

**DOI:** 10.1101/2021.10.03.462951

**Authors:** Prabhjot S. Mundi, Filemon S. Dela Cruz, Adina Grunn, Daniel Diolaiti, Audrey Mauguen, Allison R. Rainey, Kristina C. Guillan, Armaan Siddiquee, Daoqi You, Ronald Realubit, Charles Karan, Michael V. Ortiz, Melissa Accordino, Suzanne Mistretta, Frances Brogan, Jeffrey N. Bruce, Cristina I. Caescu, Richard Carvajal, Katherine Crew, Guarionex Decastro, Mark Heaney, Brian Henick, Dawn Hershman, June Hou, Fabio Iwamoto, Joseph Jurcic, Ravi P. Kiran, Michael Kluger, Teri Kreisl, Nicole Lamanna, Andrew Lassman, Emerson Lim, Gulam A. Manji, Guy McKhann, James McKiernan, Alfred Neugut, Kenneth Olive, Todd Rosenblat, Gary K. Schwartz, Catherine Shu, Michael Sisti, Ana Tergas, Reena Vattakalam, Mary Welch, Sven Wenske, Jason D. Wright, Hanina Hibshoosh, Kevin Kalinsky, Mahalaxmi Aburi, Peter A. Sims, Mariano J. Alvarez, Andrew L. Kung, Andrea Califano

**Author notes:** These authors contributed equivalently (Prabhjot S. Mundi, Filemon S. Dela Cruz).

## Abstract

Predicting tumor sensitivity to antineoplastics remains an elusive challenge, with no methods demonstrating predictive power. Joint analysis of tumors—from patients with distinct malignancies who had progressed on multiple lines of therapy—and drug perturbation transcriptional profiles predicted sensitivity to 28 of 350 drugs, 26 of which (93%) were confirmed in low-passage, patient-derived xenograft (PDX) models. Drugs were prioritized based on their ability to either invert the activity of individual Master Regulator proteins, with available high-affinity inhibitors, or of the modules they comprise (Tumor-Checkpoints), based on *de novo* mechanism of action analysis. Of 138 PDX mice enrolled in 16 single and 18 multi-protein treatment arms, a disease control rate (DCR) of 68% and 91 %, and an objective response rate (ORR) of 12% and 17%, were achieved respectively, compared to 6% and 0% in the negative controls arm, with multi-protein drugs achieving significantly more durable responses. Thus, these approaches may effectively complement and expand current precision oncology approaches, as also illustrated by a case study.

## (A) Introduction

The ultimate objective of precision cancer medicine (PCM) is to leverage molecular-level properties of a tumor—such as gene expression, epigenetic modification, proteomics, and mutational profiles, among others—to predict sensitivity to a broad range of available therapeutic agents or to guide development of novel drugs. When predictions are conserved across a significant fraction of cancer patients, successful application of PCM principles may help generate high-likelihood hypotheses for randomized clinical trials [1, 2], and may even help prioritize candidate treatment options at the individual patient level *(personalized medicine).* Systematic application of the current PCM paradigm is largely predicated on two complementary approaches. The first one *(oncogene addiction)* is aimed at identifying *targeted therapies* based on the presence of activating mutations inducing aberrant activity in druggable oncoproteins [3]; the second *(immunotherapy)* is predicated on the discovery that specific tumor-initiated immunosuppressive programs can be abrogated by pharmacological targeting of immune checkpoints of the innate and antigen-specific host response [4]. Unfortunately, not only have these approaches shown limitations that prevent their application to the majority of cancer patients but, with some exceptions, predicting patient response to either class of drugs remains challenging.

Specifically, most tumors lack actionable mutations, and even when detected their pharmacological targeting often fails to abrogate tumor viability, with benefits limited to specific cancer types or contexts. Moreover, even in patients who initially respond, the typical outcome of either approach is the relatively rapid development of drug-resistance as a result of cell-adaptive processes, inherent genetic heterogeneity, allowing for clonal selection—*i.e.,* via emergence of secondary mutations—phenotypic plasticity allowing reprograming to a drug-resistant state [5], and suppression of immunogenic neoantigen expression [6]. Consistently, multiple studies have shown that only 5% to 11% of cancer patients derive some clinical benefit, mostly transitory, from targeted therapy [7]. As such, accurate patient-level prediction of sensitivity to a vast repertoire of clinically relevant drugs remains a highly elusive problem.

Moreover, given the paucity of actionable mutations and the role that a myriad of additional genetic events may have in modulating drug sensitivity to oncogene-directed therapy, it is becoming increasingly difficult to identify large subsets of a tumor type that are sensitive to the same drug. This contributes to an ever increasing fine-grain stratification of the therapeutic landscape, prohibitively narrows selection criteria for clinical trials, and increases costs of drug development. Thus, there is an urgent need for novel methodologies for the systematic identification of more universal, non-oncogene tumor dependencies that are pharmacologically accessible and less prone to acquired resistance.

We have shown that, within individual tumor-subtypes, cancer cells can adopt only a relatively limited, discrete, and remarkably stable repertoire of transcriptional states [8]. These states are mechanistically controlled by tightly autoregulated Tumor-Checkpoint (TC) modules, comprising small, yet highly conserved sets of Master Regulator (MR) proteins, responsible for canalizing the effect of mutations in their upstream pathways [8–10]. We have also shown that TC-modules are highly enriched in MRs and MR-pairs, whose genetic [11–13] or pharmacological [14–16] inhibition can collapse the entire TC-module activity, thus abrogating tumor viability *in vitro* and *in vivo*. As such, MRs and the TC-modules they comprise are emerging as a novel, actionable class of tumorspecific, non-oncogene dependencies. Interestingly, MR and TC-module conservation is even more evident at the single cell level, where distinct, transcriptionally stable states controlled by equally distinct TC-modules are not commingled [17, 18].

An important observation is that we have shown that virtually identical TC-modules induce the same transcriptional state in tumors with highly divergent mutational profiles [8]. As such, TC-modules may represent largely mutation-agnostic, and therefore more universal dependencies within a specific tumor subtype, thus providing an appealing novel repertoire of therapeutic targets to complement targeted and immune therapy. Moreover, since TC-modules effectively canalize the effect of entire mutational landscapes in their upstream pathways, their successful pharmacological targeting should be, at least in theory, less prone to cell adaptation and clonal selection processes, thus providing more durable responses.

In this manuscript, we test two approaches to leverage this conceptual regulatory architecture for therapeutic purposes, by targeting either a single candidate MR with a high-affinity inhibitor (OncoTarget) or an entire TC-module, as determined by the Mechanism of Action (MoA) of ~350 clinically-relevant compounds inferred *de novo* from drug perturbation profiles (OncoTreat). These approaches are predicated on the ability to accurately measure the activity of ~6,200 regulatory and signaling proteins—as defined in Gene Ontology [19], see STAR Methods—from RNASeq profiles, using the VIPER algorithm [20], which we have recently shown to compare favorably with antibody based protein measurements [21]. Specifically, OncoTarget uses VIPER to identify the most aberrantly activated proteins (*i.e.*, candidate MRs) for which a high-affinity inhibitor drug is available, thus representing a straightforward, mutation-agnostic extension of the *oncogene addiction* paradigm. The rationale is that aberrant protein activity can result not only by activating mutations in the encoding gene but also by mutations and signals in upstream pathways. In contrast, OncoTreat leverages large-scale RNASeq profiles from patient-matched cell lines perturbed with clinically relevant compounds to experimentally assess their ability to invert the activity of entire TC-modules [14].

To rigorously assess the efficacy of these two methods in prioritizing effective treatments for clinical translation, we designed a tumor-agnostic study that enrolled patients with advanced disease across 14 distinct aggressive human malignancies (the *N of 1 study* at Columbia University, IRB-AAAA7562). All subjects must have progressed on at least one standard of care systemic therapy, with the majority having received ≥3 lines of treatment and fresh tissue must be available from standard of care biopsies or resections, resulting in *N* = 117 subjects enrolled to the study. Fresh tissue was implanted in immunocompromised mice to create low-passage, patient-matched PDX models for drug validation. Engrafted PDX mice were randomized to treatment with OncoTreat and/or OncoTarget predicted drugs, as well as vehicle control and suitable negative control drugs. Tumor growth was assessed by volumetric measurements. Pharmacodynamic assays, at an early time point, were also performed, to assess recapitulation *in vivo* of drug MoA predicted from drug perturbations *in vitro*. Overall analysis of PDX mice enrolled in 18 OncoTreat-based, 16 OncoTarget-based, and 13 negative control treatment arms demonstrated the effectiveness of the approach, achieving dramatically improved DCR and ORR, compared to negative controls. OncoTreat and OncoTarget have received NY and CA Dpt. of Health approval and are CLIA compliant [22], including for the analysis of archival *(i.e.,* FFPE) samples, thus paving the way to their clinical utilization, as we illustrate through an ultra-rare tumor case study.

Critically, stratification of predicted drug sensitivities across entire tumor cohorts shows that patients co-segregate within a small number of clusters (pharmacotypes) predicted to be sensitive to the same drugs, thus supporting use of these methodologies for the high-throughput generation of mechanism-based hypotheses for clinical trials. Finally, while both approaches dramatically outperformed treatment with negative control drugs and Vehicle control, the study confirmed that mice treated with TC-module-targeting drugs had statistically significantly more durable responses than those treated by targeting individual MR proteins. This is consistent with the expectation that TC-modules provide more universal targets that are less amenable to adaptation and selection mechanisms.

## Results

To translate the ability to accurately measure protein activity in individual tumor samples [20] and even in single cells [21, 23] to the clinics, we have developed two complementary, RNA-based CLIA-compliant and New York State and California Dpt. of Health approved assays. The first one, *OncoTarget,* identifies actionable, aberrantly activated proteins (p ≤ 10^-5^, as measured by VIPER), for which a high-affinity inhibitor is available. To assess target actionability, we analyzed DrugBank [24], the SelleckChem database [25], published literature, and public information from drug development pipelines, resulting in a curated list of 180 proteins representing validated, high-affinity targets of clinically-relevant small molecule compounds (**Table S1**). Conceptually, OncoTarget represents a tumor-type-agnostic, mutation-independent reformulation of the oncogene addiction paradigm [3], which extends the approach beyond established mutated oncoproteins to any actionable, aberrantly activated protein. This includes proteins that are rarely if ever mutated in cancer, such as topoisomerases, chromatin remodeling enzymes, and proteins aberrantly activated by autocrine, paracrine, or endocrine signals (**Figures 1 and S1A-B**).

**Figure 1.**
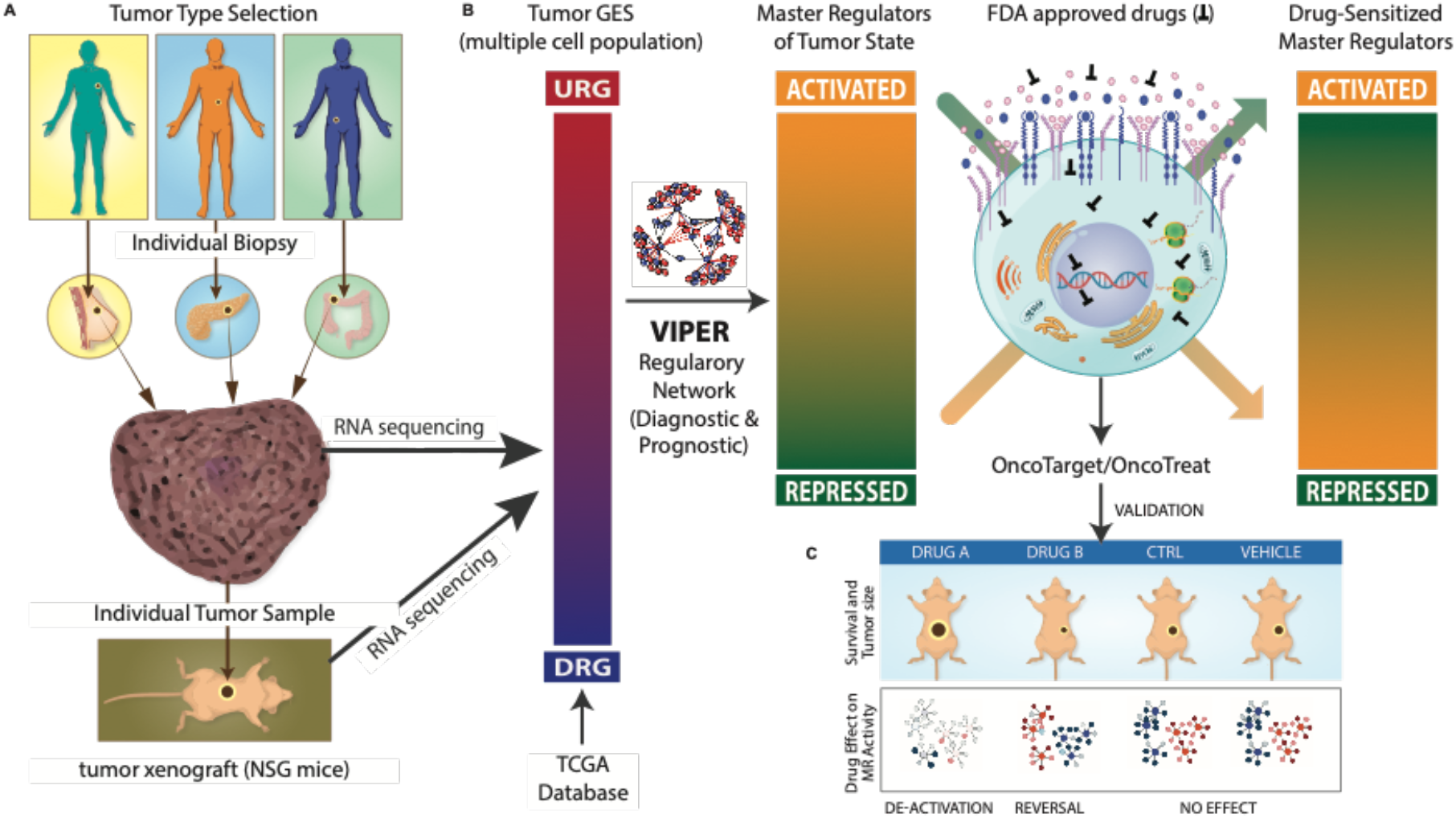
N of 1 preclinical trial conceptual diagram. **(A)** Adults with metastatic solid tumors with progression or intolerance to all standard treatments and with accessible site for biopsy are enrolled. Fresh tumor tissue from biopsy is partitioned for (*i*) clinical histopathology review, (*ii*) xenografting into immunodeficient mice, and *(iii)* mRNA profiling (RNASeq). If engraftment is successful, the mature P0 passage tumor is also profiled by RNASeq to confirm candidate MR conservation between patient tumor and PDX (OncoMatch). **(B)** Use of VIPER, OncoTarget, and OncoTreat analysis to predict optimal drugs for PDX treatment. (i) mRNA profiles are generated from tumor samples. (ii) A gene expression signature (GES) is generated by comparing the tumor profile to a large pan-cancer RNASeq compendium (reference) comprising all TCGA samples. (iii) Cancer-type specific network(s) are used to interrogate the GES to identify the most aberrantly activated and inactivated proteins (*i.e.*, candidate MRs) by VIPER analysis. *(iv)* OncoTarget identifies the most aberrantly activated proteins among those for which a high-affinity inhibitor drug is available (*i.e.*, druggable MRs)—*e.g.,* receptor and intracellular kinases, cell surface molecules, and enzymes involved in epigenetic regulation. (*v*) OncoTreat identifies the drugs inducing the strongest activity inversion of all candidate MRs (*i.e.*, TC-module inverter drugs) by VIPER analysis of drug perturbation profiles generated by treating context-relevant cell line models with available approved and experimental [antineoplastic] drugs. **(C)** Candidate drugs are prioritized based on prediction *p*-value, conservation of prediction based on the PDX RNASeq profile, and clinical relevance. Mice from the P1 passage are randomized into candidate drug arms, a negative control drug arm, and a vehicle control arm.

The second one, *OncoTreat,* leverages RNASeq profiles of representative cell lines, treated with a large repertoire of antineoplastic agents, to identify TC-module-inverter compounds that statistically significantly (p ≤ 10^-5^) invert the activity of the top 50 candidate MR proteins *(i.e.,* 25 most activated and 25 most inactivated), as assessed by VIPER analysis (**Figures 1 and S1C**) [14]. We use 50 MRs for two reasons: first, a fixed MR number is necessary to make the statistics of their activity inversion comparable across samples (see STAR methods); second, because we recently showed that, on average, across 25 cancer cohorts, ≤ 50 MRs are necessary to account for the integration of mutational events in their upstream pathways, at the individual sample level [8].

Critically, OncoTreat predicts TC-module-inverter drugs with no *a priori* knowledge of their MoA. Indeed, MoA is elucidated *de novo,* in proteome-wide fashion, by measuring the differential activity of regulatory and signaling proteins in drug vs. Vehicle control-treated cells, see STAR methods for an in-depth description.

### Model Fidelity assessment

A critical OncoTreat requirement is the availability of a comprehensive repertoire of gene expression profiles representing the transcriptional state of a tumor tissue, following treatment with a repertoire of antineoplastic compounds of interest, as well as Vehicle control (DMSO). To maximize the statistical power of the MR activity inversion analysis, these assays must be conducted in *high-fidelity* cell lines, pre-selected based on their ability to recapitulate the patient tumor’s MR activity profile. Similarly, validation should be performed in PDX models that also represent high-fidelity models for the tumor of interest.

Cell line and PDX model fidelity to human tumors was assessed based on the normalized enrichment score (NES) of the top 50 patient-specific activated and inactivated MRs in differentially activate and inactive proteins in each cell line or PDX, respectively, by analytic-rank based enrichment analysis [20] (aREA: p ≤ 10^-10^, Benjamini–Hochberg, BH-corrected). To generate a differential expression signature for VIPER-based protein activity analysis of patient and PDX tumors, we compared each sample to a common reference represented by the entire TCGA repository. To obtain an equivalent metric for cell lines, which would not be biased by their higher proliferative nature, we compared each cell line against a large cancer cell line repository— including the Cancer Cell Line Encyclopedia (CCLE) [26] and the Genentech Cell Line Screening Initiative (gCSI) [27].

In **Figure 2A**, we show the fidelity of the top 12 matching breast cancer cell lines to basal breast cancer tumors in the TCGA repository; BT20 emerged as one of the top five candidates based on the number of tumors (78 of 173), whose top 50 MRs are conserved at significance threshold *p* ≤ 10^-10^. In **Figure 2B**, we show the fidelity of four patient-matched cell lines to five patients in the study for which OncoTreat-based drugs were predicted, as well as the match between ASPC1 and a pancreatic cancer tumor, even though drug perturbation profiles in ASPC1 cells were not completed in time for *the vivo* validation studies, see additional MR-level characterization (**Figure S3**). Specifically, the BT20 cell line emerged as an excellent model for tumor BC-32398 (normalized enrichment score, NES = 14.5, *p* = 10^-48^), a very good model for BC-97359 (NES 8.0, *p* = 10^-15^), but a poor model for BC-50291 (NES −3.9, *p* = 1); both GIST cell lines GIST430 and GISTT1 were excellent matches for GIST-81050 (*p* < 10^-40^) despite not harboring the patient *SDHB*^Del^/*KRAS^G12D^* alterations, but rather canonical *KIT* mutations; IOMM was a weaker but statistically significant match for CNS-16474 (NES 3.3, *p* = 0.0005).

**Figure 2.**
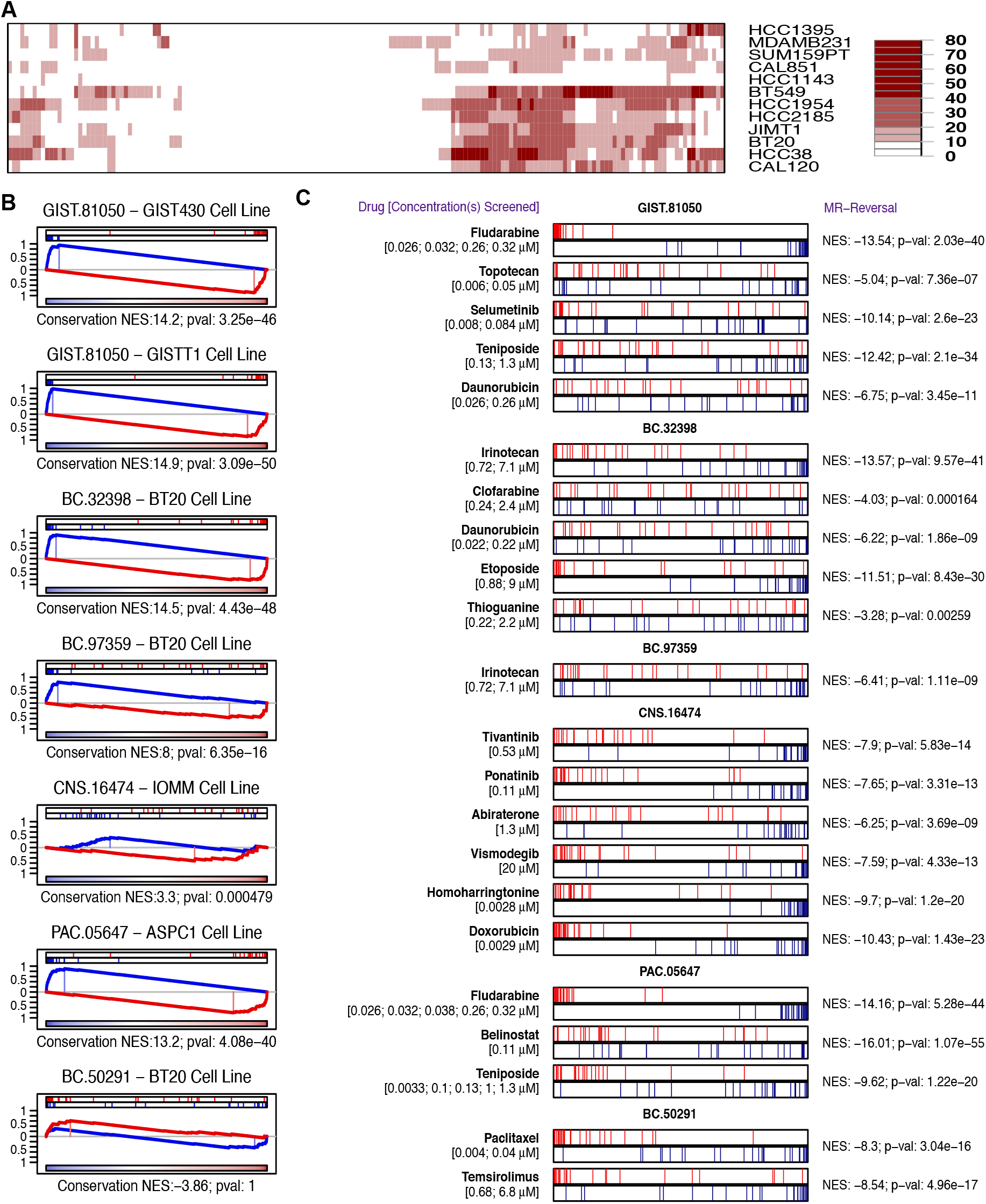
**(A)** High throughput drugs screens have been completed in over two dozen cancer cell lines that are most representative of patient tumor cohorts (PanACEA database). An example is shown, with BT20 emerging as one of the top five out of 97 breast cancer cell line profiles (top 12 shown in heatmap, corresponding to -Log_10_ (p-value) of the fidelity match), for recapitulating TC-module activity in the TCGA basal breast cancer cohort. **(B)** The enrichment of tumor-specific candidate MRs (*i.e.*, collectively the TC-module) in proteins differentially active in basal cell line profiles, is assessed through directional enrichment analysis (aREA). For four of the six tumors for which OncoT reat predictions were made, activated and inactivated candidate MRs were highly enriched in the most activated and inactivated proteins of the matched cell line model. Normalized Enrichment Scores (NES) were in the range of 8.0 to 14.9, with associated *p*-values meeting the pre-defined match threshold *p* ≤ 10^-10^. For the meningioma case (CNS.16474), the match was weaker, albeit still statistically significant (*p* = 0.0005). Finally, for the third breast cancer case (BC-50291), BT20 was not statistically significant in terms of MR recapitulation. **(C)** OncoTreat based assessment of TC-module MR activity inversion—*i.e.*, top 25 (red lines) and bottom 25 (blue lines) patient-specific MRs. For all OncoTreat-predicted drugs selected for *in vivo* validation, strong MR activity inversion (p ≤ 10^-5^) was detected following drug treatment of the patient-matched cell line, with inactivation and activation of positive and negative MRs, respectively. For the pancreatic tumor case (PAC.05647), drug perturbation profiles in the ASPC1 cell line became available only recently, thus OncoTreat analysis was based on integrated drug MoA signatures from the available cell lines, BT20, GIST430, GISTT1, and IOMM by Fisher’s method.

A critical point is that the purpose of patient-fidelity assessment is to predict and validate drug MoA in tissue contexts that optimally recapitulate the MR activity-profile of the target tumor. As a result, patient-matched cell lines are not required to also recapitulate other biologically relevant parameters, such as the mutational profile and histology of the tumor. Indeed, we have shown that MR conservation is a sufficient criterion to predict tumor-relevant drug MoA, as assessed from drug perturbation profiles, with predictions validated in both patient-derived primary cells and patient-derived explants with high statistical significance [28].

### Perturbation Profile Generation

We generated RNASeq profiles in the 4 patient-matched cell lines at 24 hours—as well as at 6h in BT20, GISTT1 and GIST430 to assess early response— following perturbation with each compound at two sublethal concentrations, the 48-hour EC_20_ and one tenth of this concentration, as determined by 10-point dose response curves. These highest sublethal concentrations were selected to focus the analysis on drug MoA rather than non-specific effector proteins of cells undergoing significant stress or death processes [29, 30]. To avoid testing drugs at non-physiologically relevant concentrations, we also capped tested concentrations at the drug’s *C*_Max_, defined as the maximum tolerated serum concentration from published human studies.

DMSO was selected as a universal solvent and Vehicle control. Multiplexed, low depth (1 to 2M read) RNASeq profiles were generated using 96-well plates via the PLATESeq methodology, using fully automated microfluidics for increased throughput and reproducibility [31]. Drug MoA— defined as the drug-mediated differential activity of all regulatory and signaling proteins—was assessed by VIPER analysis of each drug-treated sample vs. eight DMSO-treated controls included in each plate to avoid plate-dependent batch effects.

In summary, each drug perturbation profile was used to rank regulatory and signaling proteins from the most inhibited to the most activated following treatment with a specific drug at its maximum sublethal concentration vs. Vehicle control. On average, ~350 drugs were profiled in each cell line, including 138 FDA approved antineoplastics, ~170 late-stage experimental drugs in cancer clinical trials, as well as a variable number of additional compounds from diverse libraries, with cell line-specific EC50 ≤ 10 uM (**Table S2**). Since study initiation, we have now generated a Pancancer Activity by Enrichment Analysis database (PANACEA), comprising the drug MoA profiles of ~350 compounds whose activity was profiled in 23 patient-matched cell lines, representing 15 malignancies (**Table S3**). Access to the full PANACEA resource will be made available via a companion publication.

### Interactome Generation

To measure MR activity, VIPER requires a comprehensive molecular interaction dataset *(interactome)* representing the tumor context-specific transcriptional targets *(regulon)* of each protein. For these analyses—including protein activity measurement in patient, PDX, and cell line related samples—interactomes were generated by ARACNe analysis [32, 33] of RNASeq profiles from tumor-matched cohorts with ≥ 100 samples (**Table S4**).

### Study protocol and rationale

To systematically benchmark the OncoTreat and OncoTarget tests, we designed an innovative clinical study, with a preclinical endpoint based on *in vivo* tumor volume measurements following treatment with predicted drugs in low-passage PDX models. The goal was to both evaluate the efficacy of computationally predicted drugs and to assess the feasibility of the approach in the clinic (**Figures 1 and S2**). The N of 1 study enrolled 117 patients with advanced malignancies that were refractory or intolerant to standard of care treatment, representing over 20 unique cancer subtypes, including several rare and orphan ones (**Table S5**). Eligible subjects were required to have an anticipated life expectancy of at least six months, as determined by the treating oncologist. Clinically indicated biopsies or tumor resections were performed at the request of their treating oncologist; consent to allow a portion of the fresh specimen to be processed for RNASeq profiling and transplant into immunodeficient mice was required. A total of 84 implantations were possible, based on tissue availability and histology, leading to successful engrafting of 39 PDX models (46.4% take rate). Nineteen of the 39 PDX models were passaged at least once, and mature P0 passages were characterized by RNASeq and VIPER analyses to assess fidelity (**Table S5**).

### *In vivo* validation strategy

To achieve rigorous validation of OncoTreat and OncoTarget predicted drugs, we relied on the first seven PDX models that successfully engrafted and could be passaged to P1 for drug testing. These include three triple negative breast cancers (BC-32398, BC-97359, BC-50291), a pancreatic ductal carcinoma (PAC-05647), a colon adenocarcinoma (CAR-23659), a *KIT^WT^/PDGFR^WT^* gastrointestinal stromal tumor harboring *KRAS^G12D^* and a germline *SDHB* deletion (GIST-81050) in an adolescent/young adult, and a recurrent WHO grade II anaplastic meningioma (CNS-16474). The results of targeted genomic sequencing and clinical characteristics of the seven patients are summarized (**Table 1**), including extensive prior therapies with four of the seven subjects having received at least three lines of treatment before enrollment. Importantly, for all seven cases, targeted sequencing failed to identify an actionable alteration, as assessed by their treating physician at the time, thus negating potential targeted therapy approaches. Drugs to be validated in the PDX therapeutic study were selected by several criteria related to their antineoplastic nature, prior use in the patient, statistical significance, and other criteria (see STAR Methods), as follows: (*a*) Only drugs classified as antineoplastic agents were considered for validation, *(b)* Drugs were eliminated if the patient previously received the specific drug, (*c*) Drugs were prioritized based on their prediction p-value from OncoTreat and OncoTarget analysis of patient tumors, (*d*) OncoTreat predictions were selected over OncoTarget with comparable p-values when perturbation profiles were available from suitable models—e.g. for GIST, meningioma and breast cancer, (*e*) Drugs predicted not effective, based on direct corresponding OncoTreat or OncoTarget analysis of the PDX tumors *(i.e.,* p > 10^-5^), were eliminated, and (*f*) When multiple drugs with the same canonical MoA had similar prediction statistics, only the most clinically relevant was selected for validation.

**Table 1.** Clinical characteristics, prior systemic treatment, and tumor genomic profiling (if available).

| ID | Age at Diagnosis | Gender | Race/Ethnicity | Primary Tumor/<br>Histopathology | Stage at Diagnosis | Biopsy Site | Prior Systemic Treatments | Tumor Genomics<br>(Pathologic Variants) |
| --- | --- | --- | --- | --- | --- | --- | --- | --- |
| GIST.81050 | 17 | M | Caucasian/Persian | Gastrointestinal<br>Stromal Tumor<br>(GIST Sarcoma) | IV | Brain<br>Metastasis | Intraperitoneal mitomycin<br>Sunitinib x 1 cycle<br>Trametinib x 2 cycles<br>Regorafenib x 5 cycles<br>CB-839 (glutaminase inh) x 1 cycle *on a clinical trial<br>Pazopanib x 2 cycles | KRAS<br>CDKN2A<br>SDHB (splice site;<br>germline)<br>ZRSR2 |
| BC.32398 | 48 | F | Caucasian | Triple Negative<br>Breast IDC (ER 0,<br>PR 30%, HER2 0,<br>Ki-67 50%) | III | Peritoneal<br>Metastasis | Paclitaxel x 12 weeks + propanolol *on a clinical trial<br>Adriamycin + Cyclophosphamide x 4 cycles<br>Tamoxifen x 6 months<br>Leuprolide + Anastrozole x 3 months | PIK3CA<br>TP53 (x2 mutations)<br>ATM<br>NIPBL<br>SETD2<br>RPS6KB2<br>EPHB1 (x2 mutations)<br>FAM175A |
| CAR.23659 | 83 | F | Caucasian | Colon<br>adenocarcinoma,<br>microsatellite<br>stable | II | Liver Metastasis | Capecitabine + Oxaliplatin x 3 cycles | KRAS<br>ERBB2<br>APC<br>TP53<br>STAG1<br>RAD51C |
| BC.97359 | 57 | F | African American | Triple Negative<br>Breast IDC (ER 0,<br>PR 40%, HER2 0,<br>Ki-67 50%) | IIIA | Chest Wall<br>Recurrence | Adriamycin + Cyclophosphamide x 4 cycles<br>Paclitaxel x 12 weeks<br>Anastrozole | AKT1<br>TP53 |
| CNS.16474 | 56 | M | Caucasian | Atypical<br>meningioma<br>(WHO Grade II) | Grade II | Meningeal/Brain<br>Recurrence | Hydroxyurea (concurrent with gamma knife radiation) | Not available |
| PAC.05647 | 59 | F | Caucasian | Pancreatic<br>Adenocarcinoma | III | Liver Metastasis | 5-FU + Irinotecan + Oxaliplatin (FOLFIRINOX) x 12 cycles<br>Gemcitabine + Irinotecan + Everolimus x 6 months | Not available |
| BC.50291 | 47 | F | Caucasian | Triple Negative<br>Breast IDC (ER 0,<br>PR 10%, HER2 0<br>but amplified by<br>FISH, Ki-67 60%) | IV | Breast | Nab-paclitaxel x 5 cycles<br>Glembatumumab vedotin x 2 cycles *on a clinical trial<br>Capecitabine x 3 months<br>Trastuzumab + Pertuzumab + Nab-paclitaxel x 3 cycles<br>Carboplatin + Gemcitabine x 5 months<br>Sacituzumab Govitecan x 9 Cycles *on a clinical trial | No mutations identified<br>(Limited breast cancer<br>specific panel) |

The 18 OncoTreat-inferred and 16 OncoTarget-inferred drugs (28 individual drugs in total, 6 of which were predicted by both tests) selected for validation in the seven PDX models, based on these criteria, as well as their prediction rationale, dosing and schedule, are summarized in **Table 2**, (see **Table S6** for further details on drug prediction and selection in each model). The *in vitro* ability of the 18 OncoTreat-predicted drugs to invert activity of the patient TC-module (top/bottom 25 MRs) in the relevant cell line model is shown in **Figure 2C**.

**Table 2.**
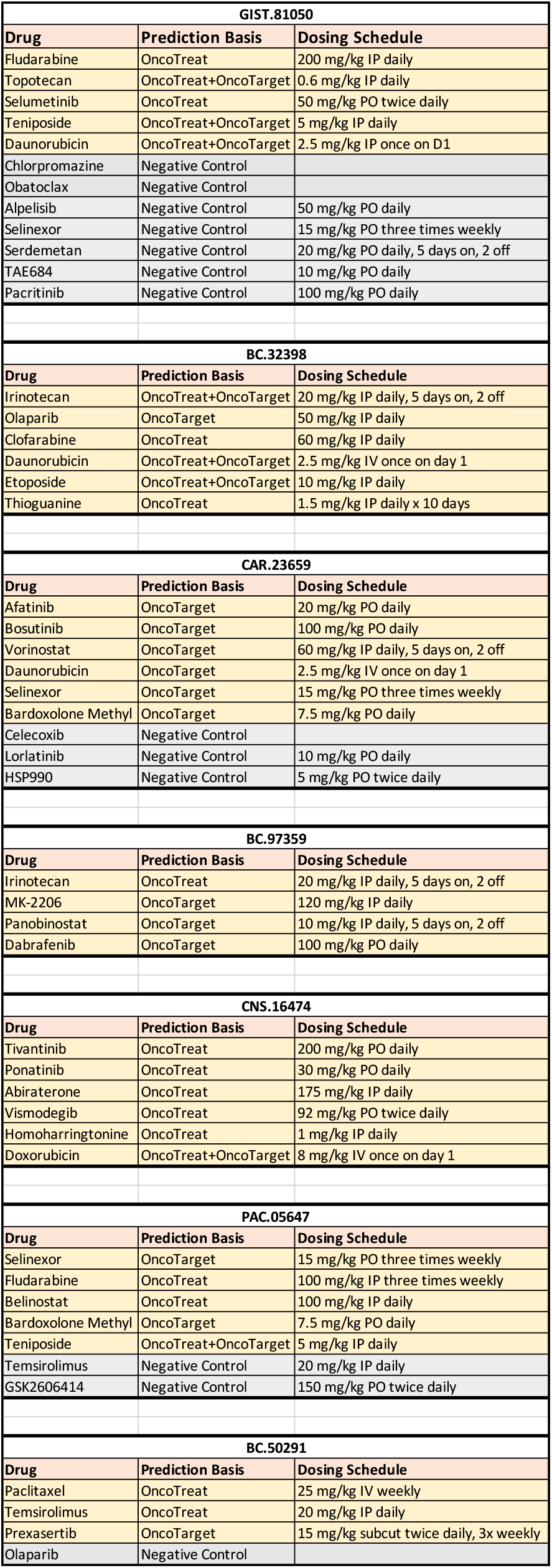
Prioritized drugs, prediction basis, and dosing schedule for PDX efficacy study.

### PDX fidelity assessment

Tumors undergo clonal evolution under various selection pressures, including available nutrients, organ-specific environment, immune editing, pharmacological treatment, and growth kinetics [34]. This is especially critical when tumors are transplanted in an immunocompromised environment, as selective pressures may be very different from those in the original human host. A critical aspect of successful treatment is to target the most common dependencies in the population of tumor cells and to efficiently identify potential resistant states that may emerge from subclonal expansion or cell adaptation. As a result, as discussed above, assessing fidelity of the PDX model tumors to the original human tumors was of paramount importance to validate drugs that would be relevant in a human context.

Several groups have described clonal drift that occurs with sequential passages in PDX models [35]. As a result, to minimize drift, we performed all therapeutic studies in the earliest passage feasible, P1 – P5. In addition, we used VIPER to assess whether the MR proteins that were the target of the drugs predicted by our analyses were conserved in the PDX tumors. Specifically, following successful engraftment and maturation of tumors (P0 passage), we performed RNASeq and subsequent VIPER, OncoTarget, and OncoTreat analyses to determine (a) the fidelity of the model in terms of Tumor Checkpoint MR activity conservation and (b) conservation of drug predictions. Drugs predicted from patient sample analysis, which were no longer statistically significant from analysis of PDX samples, were excluded from the study.

Six of the seven PDX models, GIST-81050, BC-32398, CAR-23659, BC-97359, CNS-16474, and PAC-05647 met the pre-defined match threshold (*p* ≤ 10^-10^), with analytic-rank aREA Normalized Enrichment Scores (NES) ranging from 13.8 to 17.3 (**Figures 3A and S4**). In fact, in GIST-81050 and CAR-23659 there was almost perfect conservation of patient candidate MR proteins, while in BC-32398, BC-97359, CNS-16474, and PAC-05647 there were a handful of MR proteins having different activity rank between patient and PDX. In BC-50291, however, there was no statistically significant MR activity conservation when compared to the original human tumor (*p* = 0.89).

**Figure 3.**
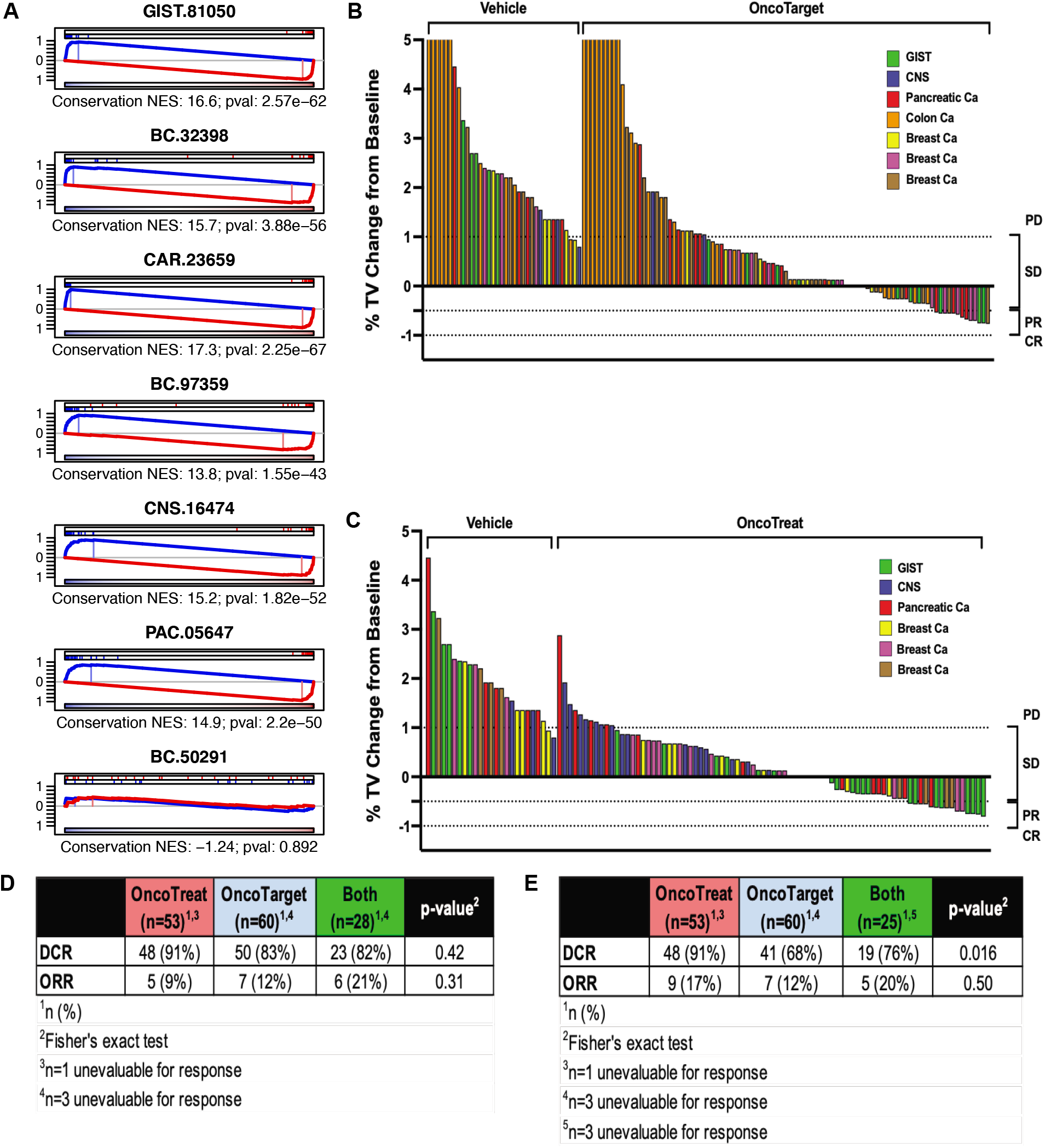
**(A)** Fidelity assessment of PDX models used for drug validation. Enrichment of tumorspecific positive and negative candidate MRs (*i.e.*, collectively the TC-module) in differentially active and inactive proteins in the P0 PDX tumor samples, assessed by analytic-rank enrichment analysis (aREA). Candidate MR activity was highly conserved in six out of seven models, but not in the (BC.50291) breast cancer model. This is potentially explained by microenvironment specific selection pressures leading to rapid PDX model drift. **(B, C)** Waterfall plots for EoS time point. Waterfall plot of mice treated with OncoTarget predicted drugs (b) in seven PDX models and mice treated with vehicle control. A disease control rate (DCR = stable disease (SD) + partial response (PR) + complete response (CR)) of 71% (n=60/85) and objective response rate (ORR = PR + CR) of 14% (n=12/85) was observed when treating with OncoTarget predicted drugs. Responses were primarily SD (n=48) and PR (n=12). Waterfall plot of mice treated with OncoTreat predicted drugs (c) in six PDX models and mice treated with vehicle control. A DCR of 86% (n=67/78) and ORR of 18% (n=14/78) were observed consisting of SD (n=53) and PR (n=14) responses. Both OncoTarget and OncoTreat were highly accurate in predicting disease control versus vehicle control (*p* < 10^-4^, by Mann-Whitney-Wilcoxon test). **(D, E)** summary of endpoints for each arm at the early TxFail time point (d) and the EoS time point (e), including a non-overlapping category for drugs predicted by both OncoTarget and OncoTreat (third column).

### PDX efficacy study

Cumulatively, significant anti-tumor responses were observed in five out of seven PDX models using drugs prioritized by either OncoTarget or OncoTreat (**Figure 3B-D and Table S7**). After expansion of PDX models for therapeutic studies, animals were enrolled onto study when tumor volumes reached 100 mm^3^. A total of 16 OncoTarget and 18 OncoTreat-predicted drugs were evaluated in individual PDX therapeutic arms (28 discrete drugs in total). Of these, six were predicted by both analyses. Response (either stable disease or partial response) was observed in 26 of 28 predicted drugs (93%) in at least one PDX mouse.

Models were treated and tumor volume measurements were recorded for at least 4 weeks (range 29 – 30 days). In order to account for differences in tumor growth rates across each model, treatment response was first evaluated on the day in which the corresponding vehicle-treated control animals met tumor volume (TV) criteria for progressive disease *(i.e.,* TV >100% increase relative to baseline), defined as the day of treatment failure (TxFail; range: 15 – 30 days, median 26 days). Significant differences in TV (relative to baseline) were observed in both OncoTreat and OncoTarget groups compared to Vehicle control (OncoTreat vs. Vehicle *p* < 0.0001 by Mann-Whitney-Wilcoxon test; OncoTarget vs. Vehicle *p* < 0.0001) with a disease control rate (DCR = stable disease (SD), partial response (PR), or complete response (CR)) of 91% (48/53 mice) and 83% (50/60), and an objective response rate (ORR = PR+CR) of 9% (5/53) and 12% (7/60) in the OncoTreat and OncoTarget cohorts, respectively (**Figure S5**). The corresponding Vehicle control cohorts for OncoTreat and OncoTarget groups showed a DCR of 12% (3/25 mice) and 11% (4/36 mice) respectively. No objective response was observed in any Vehicle control-treated mouse. Thus, at TxFail, both OncoTreat and OncoTarget-predicted drugs were highly effective, albeit with no statistically significant difference between them in DCR (*p* = 0.42, Fisher’s exact test) and ORR (p = 0.31, Fisher’s exact test). Due to unanticipated toxicity related to study treatment (tumor ulceration in a breast cancer PDX model), 3 mice were unevaluable for response in the OncoTarget cohort (BC-97359, n=2 MK-2206 arm, n=1 panobinostat arm) and 1 mouse treated with OncoTreat (BC-97359, Irinotecan arm).

We also evaluated TV at a fixed end of study (EoS) time point (range: 29 – 30 days, median 29 days), thus allowing for a longer period of response monitoring and assessment of response durability across all models. When evaluated at the EoS time point, there was a marked, statistically significant difference in OncoTreat vs. OncoTarget-predicted drug treatment response (*p* = 0.016). Indeed, DCR was 91% (48/53) and 68% (41/60), and ORR was 17% (9/53) and 12% (7/60) in the OncoTreat and OncoTarget cohorts, respectively (**Figure 3B&D**), indicating more durable responses in PDXs treated with drugs targeting the entire TC-module rather than an individual MR protein. This suggests that drug-mediated inhibition of TC-modules may implement a more effective regulatory blockade that is less likely to be circumvented by adaptive processes and clonal selection.

Similar to our prior analysis at the treatment failure timepoint, n=1 (BC-97359, Irinotecan arm), n=3 (BC-97359, MK-2206 and Panobinostat arms), and n=3 (GIST-81050, Daunorubicin and Topotecan arms) mice could not be evaluated at the arm level, due to the unanticipated toxicity.

To evaluate the specificity of drug response in the OncoTarget and OncoTreat cohorts, we identified Negative Control antineoplastic drugs for which neither OncoT arget or OncoT reat (when available) predicts an anti-tumor effect. Four PDX models (GIST-81050, CAR-23659, PAC-05647, BC-50291) were treated with 13 Negative Control drugs (**Table 2**) and Vehicle Control. At the TxFail time point (range: 7 – 30 days, median 12 days; **Table S7**), these demonstrated DCR = 53% (31/59) and ORR = 0%. However, at the EoS time point (range: 21 – 30 days, median 28 days), DCR dropped to only 6% (3/54), with an ORR of 0%. Since the TxFail time point is related to the growth rates for each model, and as Negative Control studies could not be conducted concurrently with the OncoTarget and OncoTreat cohorts, variable tumor growth rates in the models may have limited the window for response assessment as reflected by the earlier TxFail time point in Vehicle control-treated mice. Nonetheless, overall disease progression was observed across all models treated with Negative Control drugs (**Table S7**). Additionally, tumor growth inhibition (T/C% ratio) was significantly superior in the OncoTarget (mean T/C% 11%, 95% CI: −5.8 – 27.8, *p* = 8.0×10^-4^) and OncoTreat treated cohorts (mean T/C% 14.1%, 95% CI: −5.2 – 33.4, *p* = 2.0×10^-3^), compared to the Negative Control cohort (mean T/C% 48.9%, 95% CI: 29.2 – 68.5), overall *p* = 7.0×10^-4^ by ANOVA (**Figure 4C**).

**Figure 4.**
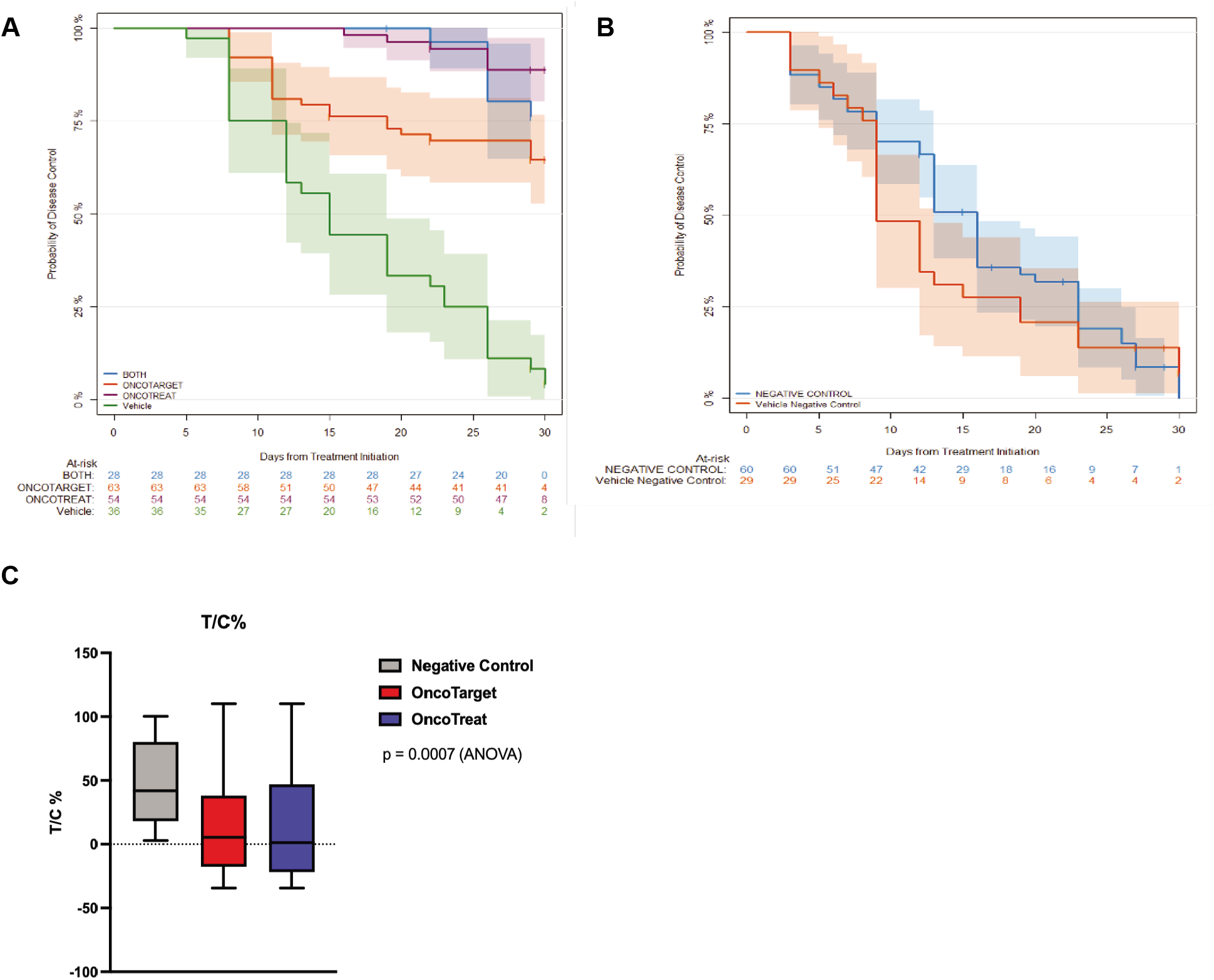
Kaplan-Meier and T/C% Tumor Growth Inhibition Analysis. **(A)** Significant disease control was observed in cohorts treated with either OncoTarget or OncoTreat predicted drugs compared to vehicle control (*p* < 10^-4^, log-rank test). There was also improved disease control with OncoTreat predicted drugs compared to OncoTarget (*p* = 0.002), but no significant difference between drugs predicted by OncoTreat alone or by both OncoTarget and OncoTreat (*p* = 0.10). **(B)** By comparison, drugs predicted to exert no anti-tumor effect based on OncoTarget or OncoTreat analyses (Negative control drugs) show no significant difference in disease control when compared to vehicle control (p = 0.38). **(C)** Boxplot of distribution of T/C% tumor growth inhibition seen in animals treated with Negative Control, OncoTarget, and OncoTreat-predicted drugs, normalized to matched-Vehicle control. There is a statistically significant difference in median T/C% in OncoTarget and OncoTreat treated mice versus Negative Control.

Cumulative Kaplan-Meier analysis for animals in the OncoTarget, OncoTreat, and OncoTreat+OncoTarget cohorts was performed (**Figure 4A**). The analysis demonstrates highly statistically significant improvement in disease control using agents predicted by either analytic approach, compared to Vehicle Control (p < 1.0×10^-4^, by log-rank test). In contrast there was no statistically significant difference between the animals in the Negative Control and concurrent Vehicle Control cohorts (*p* = 0.38) (**Figure 4B**).

Consistent with TV comparison at the EoS time point, Kaplan-Meier analysis also identified a highly significant disease control difference between the OncoTarget and OncoTreat cohorts (*p* = 2.0×10^-3^), with improved disease control in the OncoTreat cohort. Finally, no significant differences were observed between drugs predicted by OncoTreat alone versus both OncoTreat and OncoTarget (*p* = 0.10).

### Pharmacodynamic Efficacy

Pharmacodynamic (PD) studies are a critical aspect of preclinical and early clinical drug development to elucidate drug MoA and to characterize primary and acquired drug resistance. PD assessment from early, on-treatment samples helps determine whether: (a) effective, OncoTreat-predicted drugs recapitulate *in vivo* the Tumor Checkpoint MR activity inversion that occurs in cell lines *(i.e.* mechanism conservation); (b) failure of OncoTreat-predicted drugs correlates with the inability to recapitulate MR inversion *in vivo (i.e.* failure to conserve MoA), perhaps due to pharmacokinetic factors; (c) failure occurs despite MR inversion, for instance due to later cell adaptation or clonal selection; and (d) ultimately, whether assessing Tumor Checkpoint inversion in early, on-treatment biopsies may be used as a predictive biomarker of response to OncoTreat predicted drugs.

Samples for PD assessment were procured from mice sacrificed approximately three hours following the third dose, in two animals per treatment arm in five of the seven PDX models: GIST-81050, BC-32398, CAR-23659, CNS-16474, and PAC-05647 with significant conservation of patient measured Tumor Checkpoint MR activity (*i.e.* fidelity; *p* ≤ 10^-10^, **Figure 3**). Since these mice were sacrificed independent of tumor size, they were excluded from outcome assessment. OncoTreat analysis of RNASeq from drug-treated vs. Vehicle Control-treated mice was performed to assess statistical significance of MR activity inversion.

Overall, of 18 drugs predicted by OncoTreat for which PD samples were procured, all but three significantly recapitulated *in vivo* the MR-inversion (*p* ≤ 10^-5^) predicted from perturbation of MR-matched cell lines with p-values in the range of 10^-40^ (daunorubicin in BC-32398) to 10^-5^ (teniposide in PAC-05647) (**Figure 6**). Of the three drugs that failed to recapitulate the *in vitro* assessed activity, 1 was only borderline for meeting the disease control endpoint, abiraterone in CNS-16474 (**Table S7**). Belinostat also failed to recapitulate the *in vitro* MR-inversion but achieved disease control at EoS in PAC-05647. This is likely because this drug targets a chromatin remodeling enzyme and its cell state reprogramming activity may not be evident at this early time point biopsy. Finally, daunorubicin achieved disease control despite failing to achieve statistically significant MR inversion *in vivo* in the GIST-81050 model (it was however recapitulated in the BC-32398 model). However, this was due to failure to inhibit the positive MRs, while the negative MRs were effectively activated by daunorubicin (single-tail analysis *p* = 10^-2^). We have shown that negative MRs include potent tumor suppressors. As a result, their concerted activation was likely sufficient to abrogate tumor viability. In contrast, one drug (homoharringtonine) failed to achieve disease control in the CNS-16474 model, at the EoS, despite effectively recapitulating the MR-inversion predicted from the *in vitro* assays, suggesting that either clonal selection or late cell adaptation/reprogramming to a drug-resistant state may have been responsible for treatment failure.

Three of the Negative Control drugs—alpelisib, serdemetan, and pacritinib—failed to achieve significant MR-inversion activity *in vivo* and none of them achieved disease control by EoS. In contrast, TAE684 achieved significant MR-inversion activity (*p* = 10^-16^) but also failed to achieve disease control (**Figure 5**). This suggests that the early on treatment PD sample may have failed to capture the subsequently acquired cell adaptive response that ultimately led to loss of drug effect and treatment failure.

**Figure 5.**
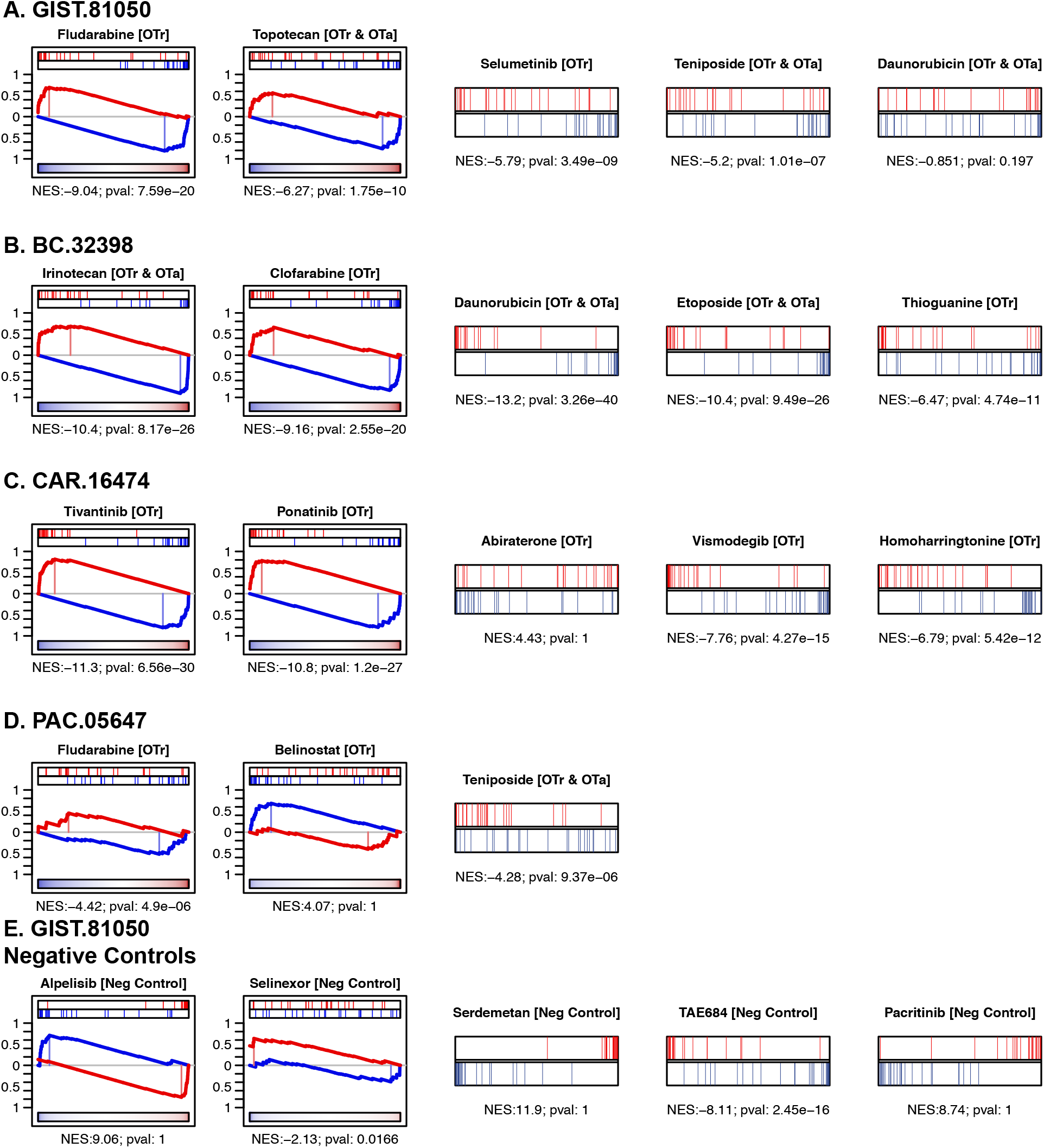
Pharmacodynamic assessment of MR activity reversal *in vivo,* in early on-treatment biopsy samples. Two mice from each drug arm were sacrificed after the 3^rd^ dose. VIPER was used to generate a differential protein activity signature from the drug-treated versus Vehicle control-treated PDX model. The enrichment of tumor-specific activated and inactivated candidate MRs (*i.e.*, collectively the TC-module), in proteins differentially inactivated and activated in the treated PDX model was assessed by analytic-rank enrichment analysis (aREA). (**A – D**) Statistically significant MR-inversion *in vivo* (*p* ≤ 10^-5^), which recapitulated the predictions from MR-matched cell lines *in vitro*, was confirmed for 15 of the 18 OncoTreat-predicted drugs. Exceptions included daunorubicin in GIST.81050, which however achieved disease control, abiraterone in CNS.16474, which induced only minimal tumor growth inhibition, and belinostat, an epigenetic modulator, in PAC.05674 which achieved disease control. **(E)** As expected, four of five negative control drugs tested in GIST.81050, did not significantly invert MR activity. TAE684 significantly inverted MR activity at this early time point. All control drugs failed to induce disease control in this model.

Taken together, 15 of 18 OncoTreat predicted drugs, including 13 of 16 that resulted in disease control, demonstrated significant *in vivo* recapitulation of *in vitro*-predicted MoA (p ≤ 10^-5^).

### Pharmacotypes identification for Clinical Trial Design

The OncoTreat and OncoTarget approaches can identify multiple candidate drugs for the treatment of the vast majority of tumors, a majority of which induced DCR or ORR when tested in the associated PDX model. Given the inherent conservation of MR protein activity within cancer subtypes, as identified by protein activity-based cluster analysis, it is not surprising to see an equivalent stratification of OncoTreat and OncoTarget predictions. Indeed, the majority of cancer types in TCGA could be stratified into 2 to 7 clusters representing patients with predicted sensitivity to the same drugs *(“pharmacotypes”)* **(Figure 6A**; Supplementary Figures Ota/Otr heatmaps). Pharmacotypes help (a) identify drugs consistently predicted across a significant fraction of patients in a specific tumor type (e.g. breast cancer) to help formulate mechanism-based hypotheses for pre-clinical and clinical trials and (b) avoid drugs that appear as uniquely relevant to a single patient, which may be false positives due to the noisy nature of RNASeq data. Consistent with the clinical observation that, as monotherapy, the vast majority of antineoplastic drugs are at best effective only in a [small] subset of patients, pharmacotypes provide a critical mechanistic link to identify patients most likely to benefit from drugs that may otherwise fail if tested broadly without a selection criterion.

**Figure 6.**
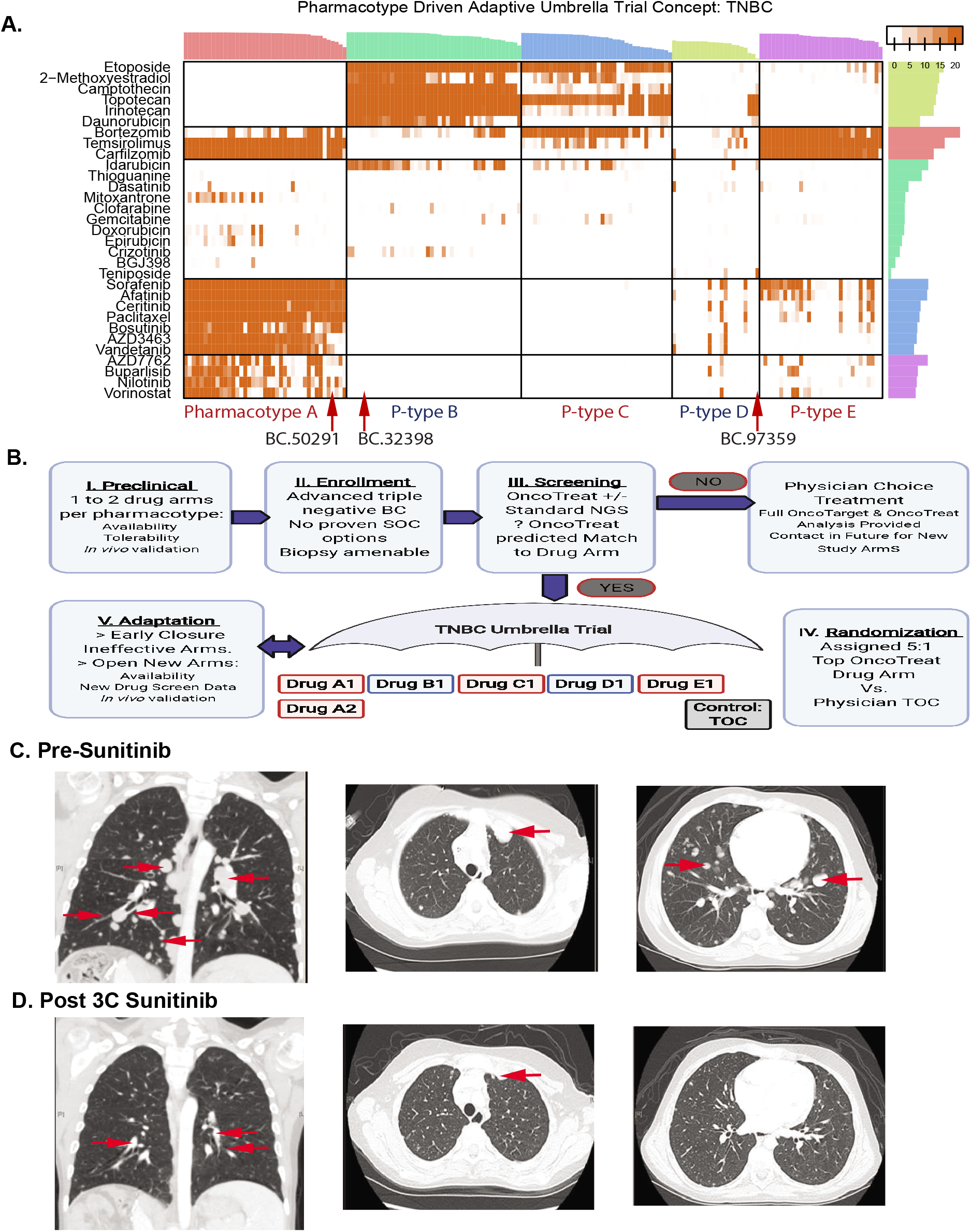
(**A**) Heatmap shows top OncoTreat predictions for 173 TCGA basal breast cancer samples, as well as for the three basal breast cancer samples from patients enrolled in the study (BC.50291, BC.32398, and BC.97359). Color code represents the –Log_10_ *p* of the prediction strength. Drug predictions cluster into five main pharmacotypes, with the three N of 1 cases clustering with samples in pharmacotype A, B, and D, respectively. (**B**) Conceptual schema for a pharmacotype-based umbrella trial concept in triple negative breast cancer. One or more drugs per pharmacotype may be prioritized as hypotheses for a follow-up umbrella clinical trial, based on availability, tolerability, and preclinical validation in relevant PDX models matching the MR profiles. Patients with advanced triple negative breast cancer who have exhausted all proven effective treatment options would be enrolled and undergo biopsy. An initial screening phase would determine if OncoTreat predicts a match to any of the open drug arms. Patients with the most statistically significant match to a specific pharmacotype would be randomized 5:1 to OncoTreat-predicted drug versus physician treatment of choice. Using an adaptive design, arms that fail to demonstrate efficacy would be closed early and new arms would be opened, as drugs become available and are validated in PDX models. (**C, D**) Response to sunitinib in a pediatric Calcifying Nested Stromal Epithelial Tumor (CNSET) patient with aberrant activation of PDGFR-B, as assessed by OncoTarget analysis. (**C**) Chest CT pre-sunitinib treatment: coronal section (top) and axial sections (middle and bottom) demonstrate numerous pulmonary metastases (red arrows) ranging from less than 1 to close to 3 cm in size. Several of the tumors were new or growing on serial scans during the preceding six months. (**D**) Chest CT following three cycles of sunitinib (six weeks each). Several of the tumors had decreased in size (red arrows) or were no longer radiologically evident. CNSET

OncoTarget, in particular, can significantly expand the pool of eligible patients for basket trials using targeted agents, while OncoTreat can be used as the basis of umbrella trials where patients with one or more cancer types can be assigned to preselected drug treatment arms based on their pharmacotype classification. To illustrate this concept, we show the pharmacotypes identified by our analysis in triple negative breast cancer (TNBC) and how this could be leveraged to design an umbrella trial (**Figure 6A-B**).

### N-of-1 Clinical Application: A Case Report

The proposed PCM framework discussed in this study is uniquely suited to identify therapeutic alternatives even for rare cancers lacking actionable mutations and standard of care options. Calcifying Nested Stromal Epithelial Tumor (CNSET) is an exceptionally uncommon primary hepatic tumor that occurs in children and young adults, with only about 40 cases reported in the literature [36]. While localized disease is often effectively cured with surgical resection, recurrent and *de novo* metastatic disease demonstrates chemotherapy resistance and there are no proven therapeutic options [37–40].

We present the case of a 14-year-old male who reported a 1-month history of abdominal pain and fatigue. A computed tomography (CT) scan of the chest and abdomen revealed a large hepatic mass with multiple satellite liver tumors and pulmonary metastases. The mass was biopsied and histopathologic evaluation was consistent with CNSET. The family was initially hesitant to initiate chemotherapy, but the patient developed progressive hepatomegaly, anorexia, weight loss, constipation, and anemia in the subsequent three months. Memorial Sloan Kettering IMPACT [41], a targeted next-generation sequencing panel covering 468 genes, was performed on a biopsy specimen and demonstrated a *CTNNB1* hotspot mutation, *TERT* promoter gain of function mutation, and *NTRK3* point mutation not known to predict response to currently available TRK inhibitors. The estimated tumor mutational burden was only 2.6/megabase, predicting a low likelihood of tumor response to immune checkpoint inhibitors.

Based on the observation that the tumor shared biological features with Wilms’ Tumor *(CTNNB1* and *TERT* mutations, WT1 and β-catenin expression by immunohistochemistry), the patient was treated with 10 neoadjuvant cycles of a “Wilms’ tumor like-regimen,” including five cycles of vincristine/irinotecan, four cycles of vincristine/dactinomycin/doxorubicin, and one cycle of cyclophosphamide/topotecan [42, 43]. There was a partial response to chemotherapy and the patient successfully underwent debulking surgery. Post-operative chemotherapy was complicated by the development of severe colitis, and the family elected to discontinue systemic therapy. Over the next six months, there was evidence of significant disease progression in the liver and lungs (**Figure 6C**), and the patient developed biliary obstruction and transaminitis that made him ineligible for clinical trials and precluded the use of most chemotherapy agents.

Given the lack of remaining viable therapeutic options, tumor tissue was sent for the CLIA-certified OncoTarget test. The most significantly activated targetable protein was PDGFR-B (*p* = 10^-7^, Bonferroni corrected). After discussing the results with the family, including the absence of clinical data on targeting PDGFR-B in this exceedingly rare malignancy, we felt that sunitinib would be the best candidate drug, given its high relative selectivity for PDGFR-B, accessibility, and safety data in the context of impaired hepatic function. The patient had a partial response to the first cycle of sunitinib (six weeks) which deepened by the end of cycle 3 (Figure 6D). Remarkably, this patient who had rapidly progressing treatment refractory cancer, has had a durable response and remains on sunitinib, now for two years from his original presentation without any appreciable side effects.

## Discussion

Approaches based on oncogene addiction [3] and immuno-oncology [4] have been both illuminating as to the potential promise that precision cancer medicine holds by driving the development of highly effective treatments for some patients, but also disappointing in their inability to benefit the majority of patients with advanced cancer in spite of large-scale efforts. We have performed extensive preclinical studies on MR proteins—establishing their role as the mechanistic determinants for integrating the effects of multiple mutations in their upstream pathways and for activating regulatory programs *(cancer hallmarks)* necessary for cancer cell survival [8–10, 44]. Indeed, MR analysis has elucidated novel mechanisms of tumorigenesis, progression, and drug sensitivity in several cancer types [10].

Here we present extensive preclinical validation for a readily applicable, scalable, and tumor-agnostic framework, predicated on an MR-based conceptualization of cancer regulation, with the potential to radically expand the PCM landscape by providing rapid prioritization of effective drugs from the currently available armamentarium. OncoTarget and OncoTreat have several practical advantages over currently used approaches. First, they are NY/CA Dpt. of Health approved and CLIA compliant, allowing rapid deployment to the clinic. Second, based on our benchmarking of over 12,000 primary tumors from TCGA and now 100s of prospectively collected samples from patients with metastatic and treatment refractory cancers, these tests can prioritize multiple candidate treatments for virtually every patient. While we certainly do not expect all identified candidates to be effective in the clinic, it is an important starting point that is simply not available to cancer patients at this time, especially for rare tumors like the one reported in our case study, which can also allow treatment prioritization based on reimbursement, toxicity, and other clinically-relevant parameters. Third, the presence of well-defined pharmacotypes—*i.e.*, tumors with shared predicted drug sensitivity, in virtually all cancer cohorts that we have studied—supports the prioritization and evaluation of predictions through standard basket and umbrella trials, respectively, with the OncoTarget and OncoTreat tests as widely applicable companion diagnostics. This includes patients across multiple malignancies predicted to share sensitivity to the same drug or drugs. Fourth, given the demonstrated role of MRs as regulatory bottlenecks, whose activity is strictly necessary for tumor state maintenance via canalization of the tumor’s genetic alteration repertoire and aberrant paracrine and endocrine signals, our data suggests that pharmacological targeting of TC-modules leads to more durable clinical responses than using therapy targeting individual proteins, thus providing a single-agent form of combination-therapy. Finally, the proposed approach has the potential of capturing longitudinal changes that are not necessarily driven by new mutations, such as metastatic progression [45] and therapy resistance, thus adapting the therapy to the dynamic nature of the tumor.

A critical aspect of this research is that both drug prioritization and validation were driven by models, including cell lines and PDXs, selected based on their ability to recapitulate the activity of patient-specific Tumor Checkpoints, thus specifically assessing for potential drift in PDX P0 tumors. An alternative strategy would have been to sample tumors in P1 animals prior to randomization to drug testing, but we felt mature tumors were more likely to be representative of patient tumors that were biopsied as opposed to early tumors undergoing kinetic growth in the PDX. Consistently, the vast majority of predicted drugs (15/18) recapitulated *in vivo* their MoA inferred from *in vitro* assays. Indeed, the PD assessment approach we describe using on-treatment biopsies has the potential to accurately predict treatment response within days as opposed to the usual requisite two to three months to determine response by radiologic criteria.

Due to the robustness of VIPER to low sequencing depth, we have recently shown its applicability to measure protein activity in single cells with accuracy comparing favorably with antibody-based measurements and recapitulating bulk measurements [21, 46]. As such, we are extending the OncoTarget and OncoTreat methodologies to the single cell level, thus allowing drug prioritization for independent subpopulations co-existing in a tumor with distinct drug sensitivities, thus potentially avoiding drug resistance before it leads to relapse.

## STAR Methods

### Generation of Gene Regulatory Networks

ARACNe (Algorithm for the Reconstruction of Accurate Cellular Networks) [32, 47]. The regulatory networks were reverse engineered by ARACNe from RNASeq profiles of human cancer tissue from The Cancer Genome Atlas (TCGA), Therapeutically Applicable Research To Generate Effective Treatments (TARGET), and a few other high quality publicly available mRNA datasets (**Table S4**). The RNASeq level 3 data were downloaded from NCI Genomics Data Commons [48]; raw counts were normalized and the variance was stabilized by fitting the dispersion to a negative-binomial distribution as implemented in the DESeq2 R-package [49]. ARACNe was run with 100 bootstrap iterations using an input set of candidate regulators including: 1,813 transcription factors (genes annotated in Gene Ontology Molecular Function database (GO)55 as GO:0003700—‘DNA-binding transcription factor activity’, or as GO:0003677—‘DNA binding’ and GO:0030528—‘Transcription regulator activity’, or as GO:0003677 and GO:0045449—‘Regulation of transcription, DNA-templated’); 969 transcriptional co-factors (a manually curated list, not overlapping with the transcritpion factor list, built upon genes annotated as GO:0003712—‘transcription coregulator activity’ or GO:0030528 or GO:0045449); and 3,370 signaling pathway related genes (annotated in GO Biological Process database as GO:0007165—‘signal transduction’ and in GO Cellular Component database as GO:0005622—‘intracellular’ or GO:0005886—‘plasma membrane’). Parameters were set to 0 DPI (Data Processing Inequality) tolerance and MI (Mutual Information) p-value threshold of 10^-8^. The mode of regulation was computed based on the correlation between TF and target gene expression.

### VIPER Analysis

VIPER (Virtual Proteomics by Enriched Regulon analysis). We have previously extensively validated the VIPER algorithm as a highly robust and specific tool for the accurate inference of regulatory protein activity in a context dependent manner [10, 16, 20]. Specifically, VIPER leverages accurate network maps of gene regulation, such as those produced by ARACNe [32, 47], to infer differential protein activity from differential gene expression signatures (GES), including those from single sample analysis. VIPER infers a protein’s regulatory activity through a probabilistic enrichment framework, assessing the enrichment of a protein’s activated and repressed transcriptional targets *(regulon)* in genes that are over and under expressed in a sample, akin to a multiplexed gene-reporter assay.

VIPER identifies the most aberrantly differentially active proteins as candidate MRs that mechanistically control a tumor’s transcriptional identity via their targets, as shown in multiple studies, see [10] for a comprehensive perspective. VIPER reproducibility is extremely high, such that Spearman correlation of activity profiles generated from RNASeq at 30M to as low as 50K read depth is *ρ*≥ 0.8 [20] even though correlation of the underlying gene expression profiles is low *ρ* ≤ 0.3. This allows effective extension of all VIPER-based methodologies and was instrumental in achieving New York State CLIA certification for OncoTarget and OncoTreat. While VIPER design is uniquely appropriate for regulatory proteins that directly control gene expression, including transcription factors, co-factors, and chromatin remodeling enzymes, we have shown that the algorithm is equally effective in monitoring activity of signaling proteins [20] and cell surface markers [46]. For instance, we have shown that VIPER-measured EGFR activity was a better predictor of EGFR inhibitor sensitivity, in a large panel of lung adenocarcinoma cell lines, than the presence of canonical activating mutations [20].

Critically, genetic or pharmacological targeting of aberrantly-activated proteins, as per VIPER analysis, has been shown to abrogate viability of multiple malignancies, ranging from prostate and breast cancer [12, 50], to leukemias and lymphomas [16, 51], to neuroblastoma, glioblastoma, and neuroendocrine tumors[11, 13, 14]. As such, we propose MRs and other VIPER-inferred aberrantly activated proteins as a new class of non-oncogene dependencies.

### OncoTarget Analysis

Through the use of DrugBank [24], the SelleckChem database [25], published literature, and publicly available information on pharmaceutical company drug development pipelines, we have curated a refined list of 180 proteins that are the validated targets of highly specific pharmacological inhibitors, either FDA approved or in clinical trials (**Table S1**). This manually curated target-drug(s) database is dominated by signaling proteins and other known oncoproteins. Pharmacological agents with narrow therapeutic indices, such as those targeting neurotransmitter signaling, ion channels, and vasoactive drugs were purposefully removed from the database as these drugs pose unique challenges for repurposing to treat cancer in the clinic.

OncoTarget inference of target protein activity is accomplished by applying VIPER to the individual tumor sample GES, computed as the gene-wise relative expression to the distribution of the expression of that gene in a pan-TCGA reference group consisting of over 12,000 tumors and followed by application of a double rank transformation.

Importantly, as VIPER reports a continuous measure of protein activity (normalized enrichment score: NES) based on the strength of enrichment of its regulon in the tumor GES, the absolute threshold that predicts sensitivity to a pharmacological inhibitor may vary between proteins. We used NES values with associated p-values *p* < 10^-5^ (BH-corrected) as a conservative empirical threshold for *in vivo* validation. Using this threshold, the average tumor in TCGA has 15 unique OncoTarget predictions, ranging from a mean of 4.5 in adrenocortical carcinoma (ACC) to 28.7 in renal clear cell carcinoma (KIRC). Tumors of the same cancer type tend to have conserved OncoTarget predictions, reflecting shared dependencies within subsets of a given cancer type (**Supplemental Figures**).

### OncoTreat Analysis

We have previously described the application of OncoTreat [14] to systematically elucidate compounds capable of reverting the activity of the repertoire of candidate MR proteins that regulate the transition to metastasis in enteropancreatic and rectal neuroendocrine tumors. We now describe a more readily applicable framework to all cancers that involves: 1) Defining tumor specific candidate MRs through the application of VIPER to a GES that computes gene-wise relative expression to the distribution of the expression of that gene in a pan-cancer reference group; 2) identifying relevant context-specific *in vitro* models based on conserved enrichment of patient tumor MR activity and generating high throughput RNASeq drug perturbation profiles in these models for *de novo* unbiased elucidation of MoA; and 3) prioritization of pharmacological agents whose effector proteins *(i.e.,* post-perturbation VIPER-measured drug activity signature) are enriched in the repertoire of patient tumor candidate MRs. Drugs are ranked by statistical significance of their induced MR activity inversion based on enrichment analysis of the drug signature (*i.e.*, MoA).

OncoTreat specifically assesses the ability of drugs to invert the activity of the 25 most aberrantly activated and 25 most inactivated proteins in a specific tumor sample. This number was selected because we have shown that, on average, across all of TCGA, the vast majority of genomic events identified on an individual sample basis can be found upstream of the top 25 proteins inferred by VIPER, thus supporting their status as putative MRs [8]. OncoTreat identified entinostat, a class I histone deacetylase inhibitor, among 105 profiled drugs, as effective at inverting the activity of MR proteins of metastatic gastroenteropancreatic neuroendocrine tumors (GEP-NETs), which was then validated to induce significant tumor volume reduction *in vivo* [14], leading to a clinical trial that is currently accruing patients (NCT03211988).

### Cell Line and PDXModel Fidelity (Match) Analysis

Model match was tested by independently assessing the conserved enrichment of upregulated MRs (top 25) of the patient-specific Tumor Checkpoint module in the positive tail of the VIPER-inferred protein activity signature of the model and the conserved enrichment of downregulated MRs (bottom 25) in the negative tail of the model signature. The two resulting significance values are combined by Fisher’s method. The analytic-rank based enrichment analysis (aREA) algorithm [20] was used to perform enrichment analysis, but any suitable algorithm could be substituted. In all cases, repeat RNASeq was performed onsite on acquired cell line models to ensure consistency with the publically available profiles used for pre-selection.

### Establishment of PDX models and therapeutic drug testing

All animals were maintained under barrier conditions and all experiments were performed in accordance with and approval of the Memorial Sloan Kettering Cancer Center (MSKCC) Institutional Animal Care and Use Committee (IACUC, protocol #16-08-011) and Columbia University Medical Center (CUMC) IACUC (AAAF5850). Patient tumor tissue was collected under the CUMC Institutional Review Board (IRB)-approved protocol AAAA7562 with written informed consent provided by the subject or legally authorized representative. Generation of PDX models was performed under the MSKCC IRB-approved protocol #17-387 and CUMC IRB protocol AAAJ5811. PDX models were established as previously described[52]. In summary, fresh tumor tissue was fragmented and implanted subcutaneously into nonobese/severe combine immunodeficiency IL2Rγ null, hypoxanthine phosphoribosyltransferase (HPRT)-null (NSGH) mice (Jackson Labs, IMSR catalog no. JAX:012480, RRID: IMSR_JAX:012480) and tumor engraftment monitored by visual and manual inspection. Engrafted tumors were measured twice weekly with calipers and drug treatment initiated when tumor volume (TV) reached ~100 mm^3^ (TV = width^2^ X ½ length). Early passage animals (Passage 1 – 5) were used for all therapeutic studies. Aligned with clinical response criteria, treatment response was categorized as complete response (CR, >95% reduction from baseline or no measurable/palpable tumor), partial response (PR, >50% reduction), stable disease (SD, <50% reduction and no more than 100% increase), or progressive disease (PD, >100% increase from baseline). Disease control rate (DCR) is defined as the sum of CR, PR and SD responses. Objective response rate (ORR) is defined as the sum of CR and PR responses. Tumor responses were assessed at time of treatment failure (TxFail), defined as the day in which at least 75% of vehicle treated tumors met criteria for PD for any given model in order to account for inter-model variability in tumor growth rates, or at the end of the therapeutic study period (EoS). The Mann-Whitney-Wilcoxon method was used to determine differences in the distribution of relative tumor volume between OncoTarget or OncoTreat cohorts and Vehicle control. Vardi’s test was used to evaluate differences in area under the curve (AUC) between treatment groups across models. T/C% was determined for OncoTarget, OncoTreat, and Negative control groups and differences evaluated using 2-way ANOVA. Comparisons of DCR and ORR across treatment groups (OncoTarget alone, OncoTreat alone, Both) were performed using a Fisher’s exact test or Pearson’s chi-squared test. Disease control was defined as the percentage of mice that did not meet criteria for progressive disease for the duration of the therapeutic study. Kaplan-Meier survival curves were compared and analyzed using the log-rank test. Statistical analysis was performed using R software (v3.5.0) or GraphPad Prism [v8.4.1 (RRID:SCR_002798)]. Statistical significance was defined as a p-value < 0.05.

## Supporting information

Supplementary Darwin Report

## Acknowledgements

This work was supported by The National Institutes of Health grants U01CA217858 *Cancer Target Discovery and Development (CTD2),* U54CA209997, and R35 CA197745 *Outstanding Investigator Award* to A. Califano; P30CA013696 *Cancer Center Support Grant,* S10OD012351 and S10OD021764 *Shared Instrumentation Programs* to Columbia University.

## Supplemental

**Figure S1:** Detailed schematic of the VIPER, OncoTarget, and OncoTreat methodologies.

**Figure S2:** Detailed schematic of the N of 1 trial, including overview of subject eligibility criteria, procurement of tumor tissue for standard of care clinical evaluation and study procedures – including xenografting and RNASeq/VIPER profiling for downstream OncoTarget and OncoTreat analyses, and therapeutic validation of predicted drugs in PDX models. Tumors are measured thrice weekly for efficacy evaluation and two mice in each arm sacrificed for early on-treatment pharmacodynamics assessment.

**Supplementary Report:** Example clinical OncoTarget and OncoTreat report for an ovarian cancer case. In OncoTreat, predictions are made based on *de novo* dissected drug MoA. As such, one should not assume that drugs targeting the same primary target or representing the same inhibitor class may be considered equivalent. In OncoTarget, predictions are made on specific druggable proteins, and drugs are prioritized based on target specificity and clinical availability and use.

**Figure S3.** The position of tumor-specific candidate MRs, *i.e.*, collectively the Tumor Checkpoint, in the VIPER inferred protein activity signature of cell lines is assessed through directional enrichment analysis (aREA). The activity of specific candidate MRs used for this analysis in the patient tumor sample and cell line are demonstrated in the heatmaps on the right.

**Figure S4.** Enrichment of tumor-specific positive and negative candidate MRs (*i.e.*, collectively the TC-module) in proteins differentially activated and inactivated based on VIPER analysis of the mature P0 PDX tumor, respectively, as assessed by analytic-rank enrichment analysis (aREA). The activity of the specific candidate MRs used in this analysis in the patient tumor sample (top) and PDX (bottom) are shown in the heatmaps on the right.

**Figure S5.** Waterfall plots for TxFail time point. (**A**) Waterfall plot of mice treated with OncoTarget predicted drugs in seven PDX models and mice treated with vehicle control. (**B**) Waterfall plot of mice treated with OncoTreat predicted drugs in six PDX models and mice treated with vehicle control. Both OncoTarget and OncoTreat were highly accurate in predicting disease control versus vehicle control.

**Table S1.** Pre-defined list of proteins with high-affinity inhibitor drugs for OncoTarget analysis. Inhibitors were identified by analysis of databases such as SelleckChem and DrugBank. Proteins with no associated drugs or drugs with narrow therapeutic indices (such as vasoactive or neurotransmitters) were removed. When multiple high-affinity inhibitors were identified, they were ranked based on target binding affinity, availability, and clinical utility/tolerability (subjective).

**Table S2.** Comprehensive list of clinically relevant drugs screened in PanACEA cell lines. The list is divided into three sections, including one with FDA approved antineoplastics, one with experimental antineoplastics in clinical trials, and one with non-antineoplastics and tool compounds. Drugs in the first two categories were generally evaluated in all cell lines screened while compounds in the last category were only evaluated when demonstrating an EC50 < 10μM.

**Table S3.** Current list of cancer cell lines with available drug perturbation RNASeq profiles, which together comprise the PANACEA resource. Automated high-throughput screens were performed in all cell lines using the PLATESeq platform.

**Table S4.** List of available interactomes (gene regulatory networks) that have been generated from RNASeq datasets of patient tumor samples using the ARACNe algorithm.

**Table S5.** Summary of enrollment criteria for the N of 1 trial, including numbers by cancer type, the attempt to develop a PDX, establishment of PDX, and RNASeq/VIPER profiling of PDX when established.

**Table S6.** Detailed summary of drugs prioritized for testing in PDX models, based on rank of OncoTarget and OncoTreat predictions by –log10 p-value.

**Table S7.** Detailed summary of therapeutic response at Vehicle TxFail time point (mean tumor volume versus baseline), organized by study group – OncoTreat, OncoTarget, or Negative Control – and then stratified by each individual drug arm. The study was underpowered for statistical analyses of therapeutic response in individual drug arms.

## Notes

### Competing Interest Statement

Andrea Califano Declaration of interests A.C. is founder, equity holder, consultant, and director of DarwinHealth Inc., a company that has licensed some of the algorithms used in this manuscript from Columbia University. M.J.A. is Chief Scientific Officer and equity holder at DarwinHealth, Inc. Patent 10,790,040, (titled: Virtual Inference of Protein Activity by Regulon Analysis), has issued on Sept. 29, 2020 related to the VIPER method. Columbia University is also an equity holder in DarwinHealth Inc.

### Summary of Updates

Acknowledgements; author list.

