## Supplementary Darwin Report for "Pre-clinical validation of an RNA-based precision oncology platform for patient-therapy alignment in a diverse set of human malignancies resistant to standard treatments"

### Example Precision OncoTarget - OncoTreat Clinical Report (Ovarian Ca)

#### OncoTarget

##### FDA-Approved Drugs Targeting Aberrantly Activated Oncoproteins

| DOTa Drug Score and Rank | Aberrantly Activated Oncoproteins | Drugs |
|---|---|---|
| 1. | PSMA2 ($4\times10^{-14}$), PSMB5 ($7\times10^{-10}$), PSMB1 ($9.1\times10^{-9}$), PSMB2 ($1.6\times10^{-7}$) | **Bortezomib.** Potent 20S proteasome inhibitor with Ki of 0.6 nM.<br>**Carfilzomib.** Irreversible proteasome inhibitor with $IC_{50}$ of <5 nM, displayed preferential *in vitro* inhibitory potency against the ChT-L activity in the $\beta$5 subunit, but little or no effect on the PGPH and T-L activities. |
| 2. | RPL3 ($1.1\times10^{-20}$) | **Homoharringtonine.** Inhibits protein synthesis by binding to the A-site cleft in the large ribosomal subunit, affecting chain elongation and preventing protein synthesis. |
| 3. | PSMA2 ($4\times10^{-14}$), PSMB5 ($7\times10^{-10}$), PSMB1 ($9.1\times10^{-9}$) | **Ixazomib.** MLN2238 inhibits the chymotrypsin-like proteolytic (5) site of the 20S proteasome with IC50 and Ki of 3.4 nM and 0.93 nM, respectively, also inhibits the caspase-like (1) and trypsin-like (2) proteolytic sites, with IC50 of 31 and 3500 nM. |
| 4. | HDAC3 ($2.8\times10^{-19}$) | **Vorinostat.** HDAC inhibitor with $IC_{50}$ of 10 nM.<br>**Belinostat.** HDAC inhibitor with $IC_{50}$ of 27 nM, with activity demonstrated in cisplatin-resistant tumors.<br>**Panobinostat.** Broad-spectrum HDAC inhibitor with $IC_{50}$ of 5 nM.<br>**Romidepsin.** HDAC1 and HDAC2 inhibitor with $IC_{50}$ of 36 nM and 47 nM, respectively. |

##### Investigational Agents Targeting Aberrantly Activated Oncoproteins

| DOTa Drug Score and Rank | Aberrantly Activated Oncoproteins | Drugs |
|---|---|---|
| 1. | CDK4 ($3.4\times10^{-15}$), CDK5 ($2.2\times10^{-9}$), CDK1 ($6.7\times10^{-9}$) | **PHA-848125.** PHA-848125 is a potent, ATP-competitive CDK inhibitor for CDK2 with $IC_{50}$ of 45 nM. It is >3-fold more selective for CDK2 than CDK1, 2, 4, 5, and 7. |
| 2. | HDAC3 ($2.8\times10^{-19}$) | **Entinostat.** Inhibits HDAC1 and HDAC3 with $IC_{50}$ of 0.51 $\mu$M and 1.7 $\mu$M.<br>**Givinostat.** Givinostat (ITF2357) is a potent HDAC inhibitor for HDAC2, HDAC1B and HDAC1A with $IC_{50}$ of 10 nM, 7.5 nM and 16 nM.<br>**Tacedinaline.** CI994 (Tacedinaline) is a selective class I HDAC inhibitor with $IC_{50}$ of 0.9, 0.9, 1.2, and >20 $\mu$M for human HDAC 1, 2, 3, and 8, respectively.<br>**Citarinostat.** Citarinostat, also known as ACY-241, is a potent, selective and orally available histone deacetylase (HDAC) inhibitor, with potential antineoplastic activity.<br>**Tucidinostat.** Tucidinostat (also known as HBI-8000, Chidamide, and CS-055) is a benzamide type inhibitor of histone deacetylase (HDAC) isoenzymes 1, 2, 3 and 10, with potential antineoplastic activity. |
| 3. | MAP2K2 ($7\times10^{-12}$), AURKB ($6.6\times10^{-9}$) | **BI-847325.** BI-847325is an orally available dual inhibitor of mitogen-activated protein kinase kinase (MEK) and Aurora kinases, with potential antineoplastic activity. |
| 4. | PSMB5 ($7\times10^{-10}$), PSMB1 ($9.1\times10^{-9}$) | **MLN9708.** MLN9708 immediately hydrolyzed to MLN2238, the biologically active form, on exposure to aqueous solutions or plasma. MLN2238 inhibits the chymotrypsin-like proteolytic (5) site of the 20S proteasome with IC50/Ki of 3.4 nM/0.93 nM, less potent to 1 and little activity to 2. |
| 5. | TXN ($3.9\times10^{-14}$) | **PX-12.** PX-12 irreversibly inhibits the redox protein thioredoxin, which has been associated with cancer and tumor growth. |
| 6. | MAP2K2 ($7\times10^{-12}$) | **Pimasertib.** Pimasertib (AS-703026) is a highly selective, potent, ATP non-competitive allosteric inhibitor of MEK1/2 with $IC_{50}$ of 5 nM-2 $\mu$M.<br>**CI-1040.** CI-1040 (PD 184352) is an ATP non-competitive MEK1/2 inhibitor with $IC_{50}$ of 17 nM, 100-fold more selective for MEK1/2 than MEK5.<br>**AZD8330.** non-ATP competitive MEK 1/2 inhibitor with $IC_{50}$ of 7 nM.<br>**Binimetinib.** ARRY-438162 is a potent inhibitor of MEK1/2 with $IC_{50}$ of 12 nM.<br>**Refametinib.** Refametinib (RDEA119, Bay 86-9766) is a potent, ATP non-competitive and highly selective inhibitor of MEK1 and MEK2 with $IC_{50}$ of 19 nM and 47 nM, respectively. |
| 7. | CDK4 ($3.4\times10^{-15}$), CDK1 ($6.7\times10^{-9}$) | **Flavopiridol.** Competes with ATP to inhibit CDKs including CDK1, CDK2, CDK4 and CDK6 with $IC_{50}$ of 40 nM. It is 7.5-fold more selective for CDK1/2/4/6 than CDK7. Flavopiridol is initially found to inhibit EGFR and PKA.<br>**Roniciclib.** Roniciclib, also known as BAY 1000394; is an orally bioavailable cyclin dependent kinase (CDK) inhibitor with potential antineoplastic activity. |

#### OncoTreat

##### Summary of Tumor CheckPoint-targeting FDA-Approved Drugs in Oncology

| Rank | Tumor CheckPoint Reversion Strength | Ranking of Tumor CheckPoint-targeting FDA-Approved Drugs |
|---|---|---|
| 1. | $1.2\times10^{-30}$ | **Bicalutamide.** Androgen receptor (AR) antagonist with $IC_{50}$ of 0.16 $\mu$M. |
| 2. | $6.2\times10^{-26}$ | **Bortezomib.** Potent 20S proteasome inhibitor with Ki of 0.6 nM. |
| 3. | $0.99\times10^{-25}$ | **Selumetinib.** MEK1 inhibitor with $IC_{50}$ of 14 nM in cell-free assays, also inhibits ERK1/2 phosphorylation with $IC_{50}$ of 10 nM, no inhibition to p38$\alpha$, MKK6, EGFR, ErbB2, ERK2, B-Raf, etc. |
| 4. | $3.9\times10^{-25}$ | **Midostaurin.** Broad spectrum protein kinase inhibitor. Inhibits conventional PKC isoforms ($\alpha$, $\beta$, $\gamma$), PDFR$\beta$, VEGFR2, Syk, PKC, Flk-1, Flt3, Cdk1/B, PKA, c-Kit, c-Fgr, c-Src, VEGFR1 and EGFR. Displays potent antitumor activity. |
| 5. | $4.2\times10^{-24}$ | **Thalidomide.** Thalidomide inhibits an E3 ubiquitin ligase, which is a CRBN-DDB1-Cul4A complex. |
| 6. | $1.1\times10^{-22}$ | **Rucaparib.** Rucaparib (AG-014699, PF-01367338) is an inhibitor of PARP with Ki of 1.4 nM for PARP1, also showing binding affinity to eight other PARP domains. |
| 7. | $5.6\times10^{-22}$ | **Anastrozole.** Anastrozole is a third-generation nonsteroidal selective aromatase inhibitor. |

##### Investigational Agents Targeting Tumor Checkpoints

| Rank | Tumor CheckPoint Reversion Strength | Ranking of Tumor CheckPoint-targeting Compounds |
|---|---|---|
| 1. | $1.2\times10^{-34}$ | **Triciribine.** Triciribine is a DNA synthesis inhibitor, also inhibits Akt and HIV-1 with $IC_{50}$ of 130 nM and 20 nM, respectively; does not inhibit PI3K/PDK1; 5000-fold less active in cells lacking adenosine kinase. |
| 2. | $5.5\times10^{-29}$ | **Refametinib.** Refametinib (RDEA119, Bay 86-9766) is a potent, ATP non-competitive and highly selective inhibitor of MEK1 and MEK2 with $IC_{50}$ of 19 nM and 47 nM, respectively. |
| 3. | $4.5\times10^{-25}$ | **Telatinib.** Inhibitor of VEGFR2/3, c-Kit and PDGFR$\beta$ with $IC_{50}$ of 6 nM/4 nM, 1 nM and 15 nM, respectively. |
| 4. | $7.2\times10^{-24}$ | **TAK-733.** MEK allosteric site inhibitor for MEK1 with $IC_{50}$ of 3.2 nM, inactive to Abl1, AKT3, c-RAF, CamK1, CDK2, c-Met, etc. |
| 5. | $1.2\times10^{-23}$ | **Pipobroman.** The mechanism of action is uncertain but pipobroman is thought to alkylate DNA leading to disruption of DNA synthesis and eventual cell death. |
| 6. | $8.8\times10^{-23}$ | **Alisertib.** Selective Aurora A and Aurora B inhibitor with $IC_{50}$ values of 1.2 and 396.5 nM, respectively. |
| 7. | $1.3\times10^{-22}$ | **Bafetinib.** Dual Bcr-Abl/Lyn inhibitor with $IC_{50}$ of 5.8 nM/19 nM, does not inhibit the phosphorylation of the T315I mutant and is less potent to PDGFR and c-Kit. |
| 8. | $1.4\times10^{-18}$ | **UNC0642.** G9a and GLP histone lysine methyltransferase inhibitor ($IC_{50}$ < 2.5 nM). |
| 9. | $7.2\times10^{-18}$ | **AZ12419304.** B-Raf serine/threonine-protein kinase (BRAF) and V600E mutation inhibitor |
| 10. | $10^{-17}$ | **Givinostat.** Givinostat (ITF2357) is a potent HDAC inhibitor for HDAC2, HDAC1B and HDAC1A with $IC_{50}$ of 10 nM, 7.5 nM and 16 nM. |
