## Supplementary figures and images for "Pre-clinical validation of an RNA-based precision oncology platform for patient-therapy alignment in a diverse set of human malignancies resistant to standard treatments"

### Table S1

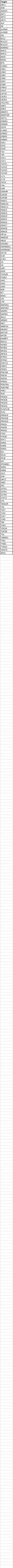

### Table S2

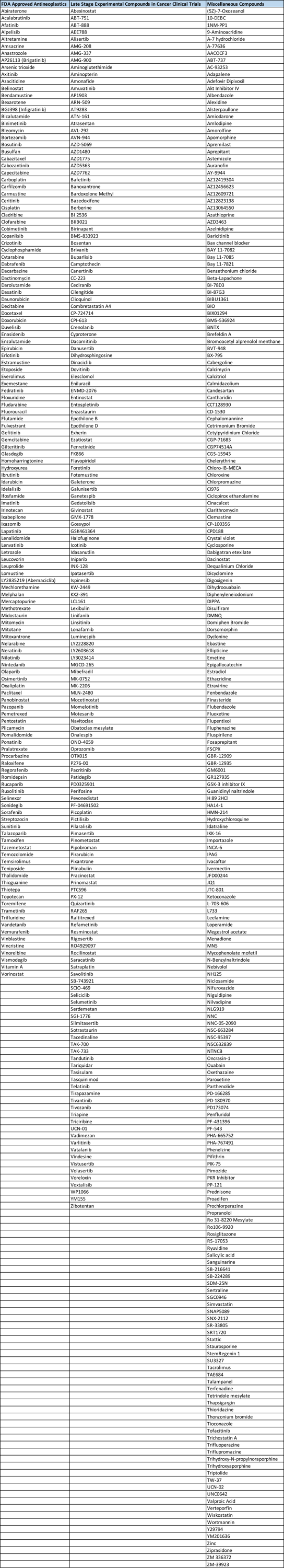

### Table S3

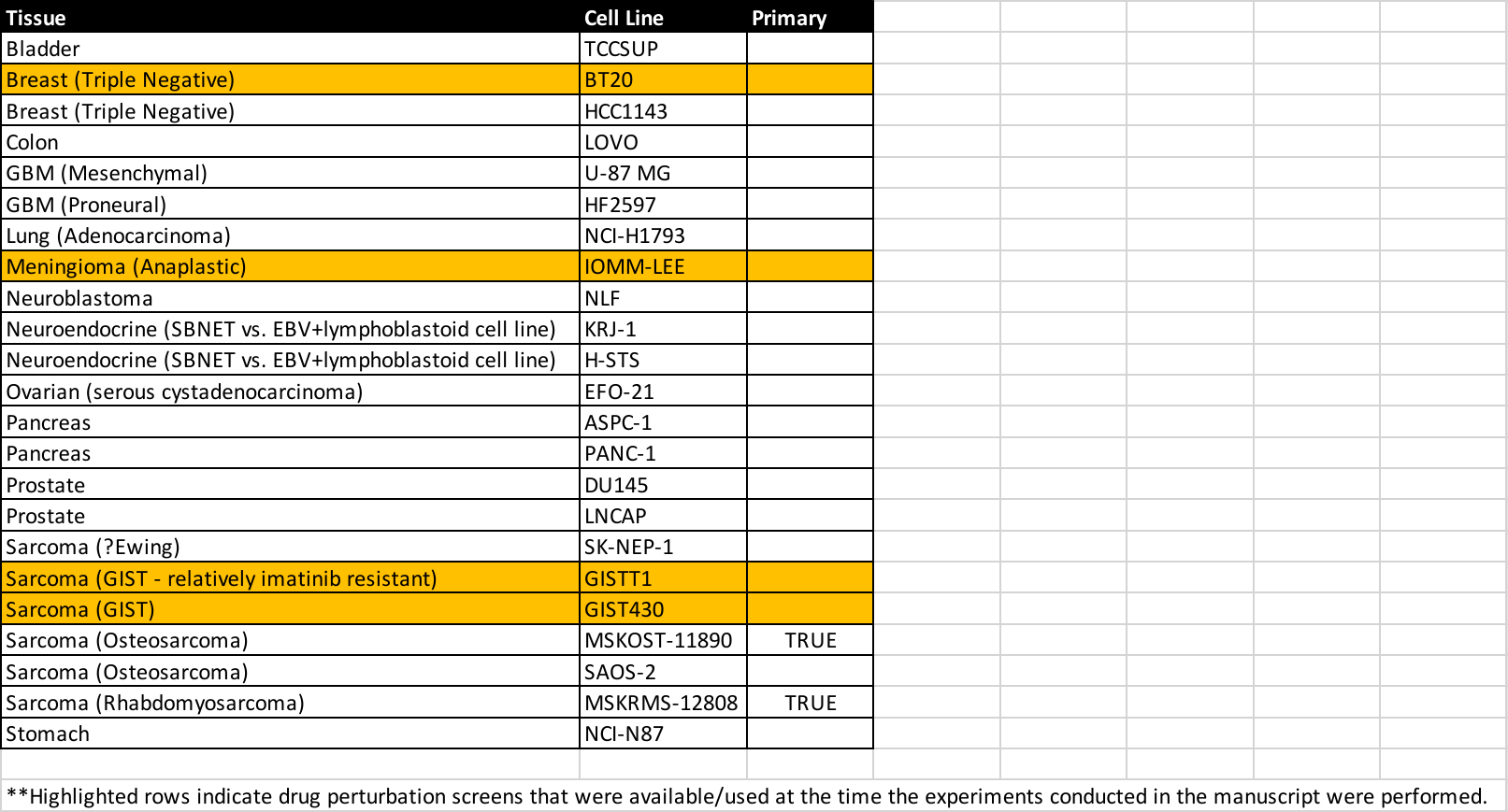

### Table S4

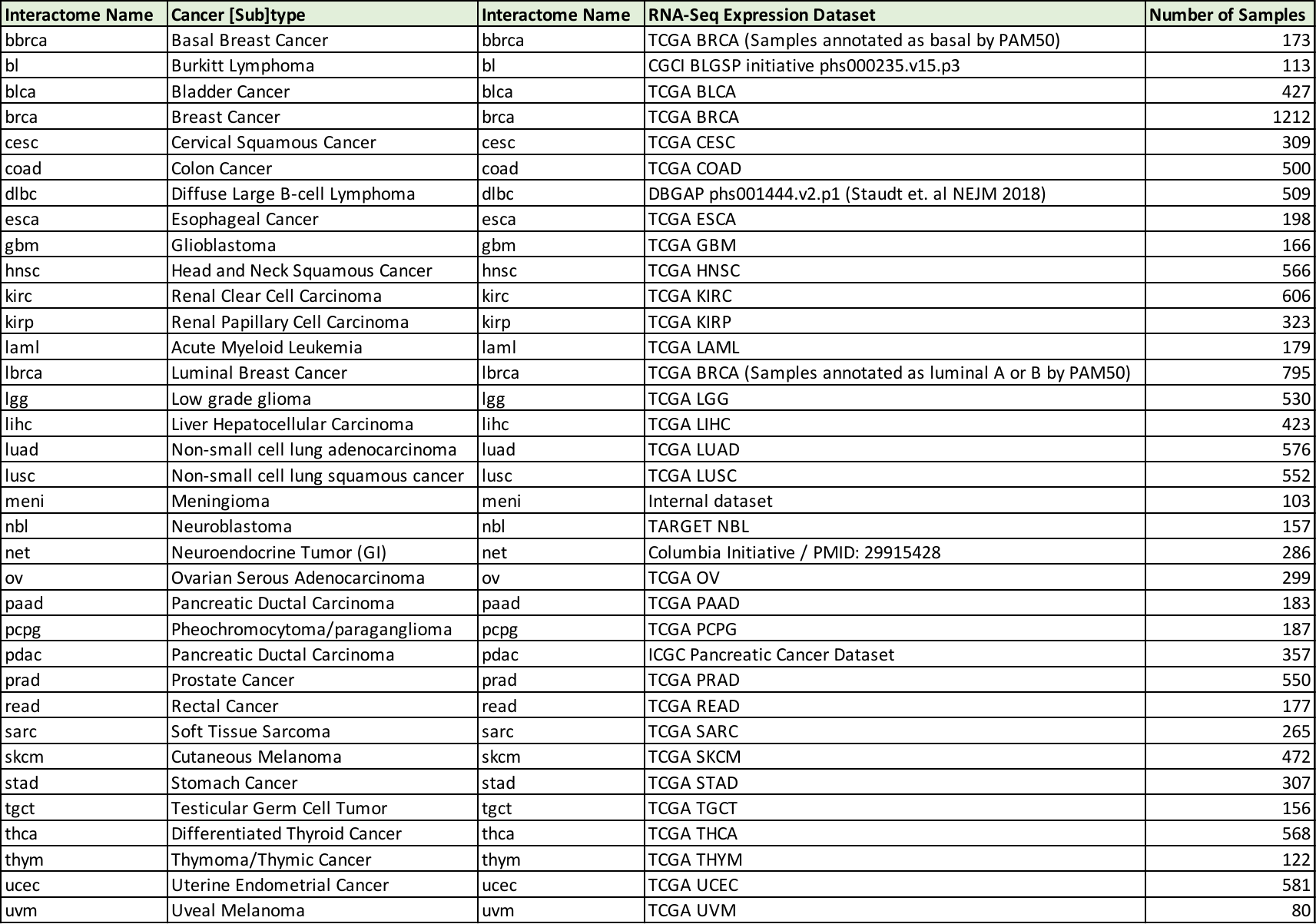

### Table S5

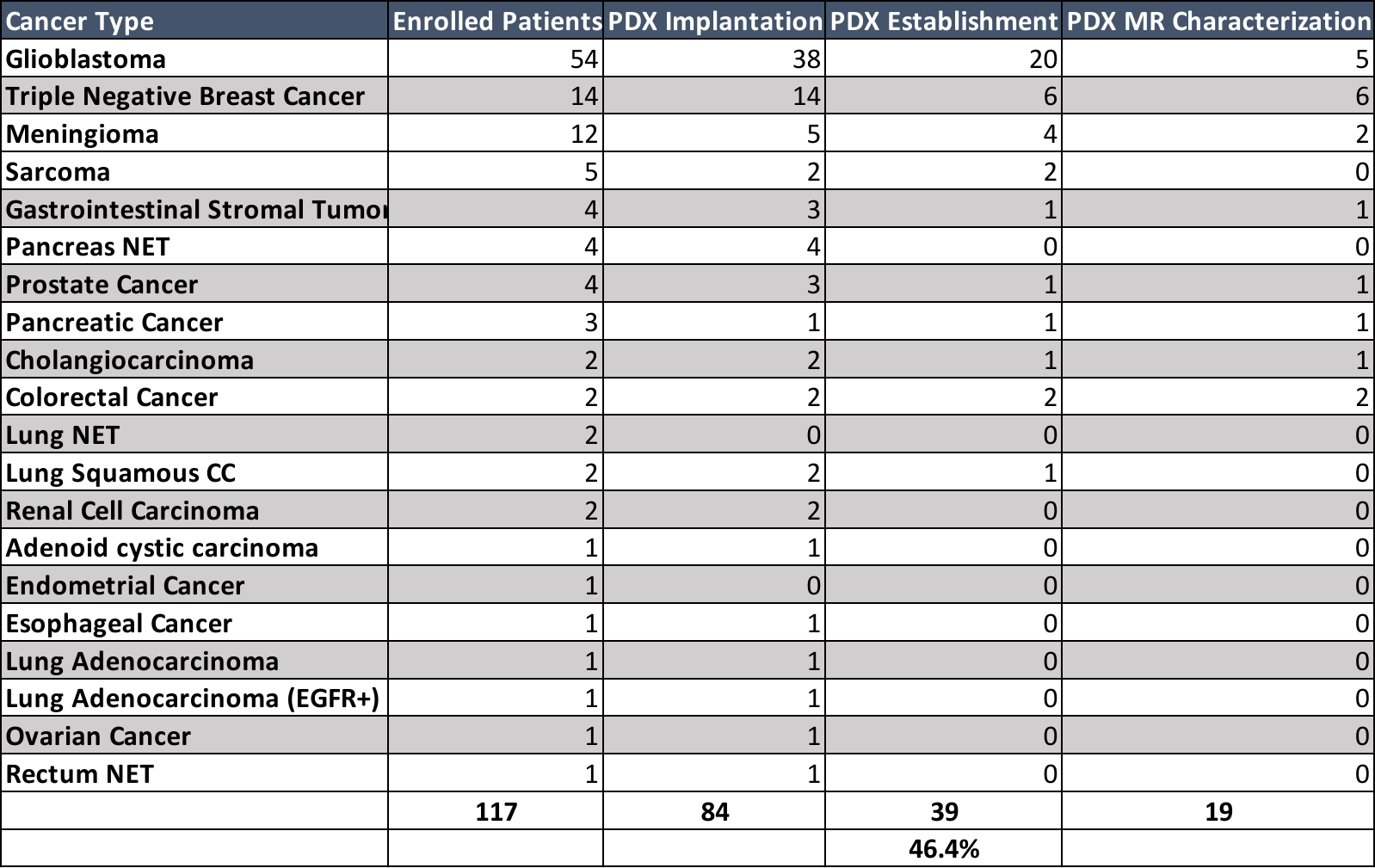

### Table S6

GIST.81050

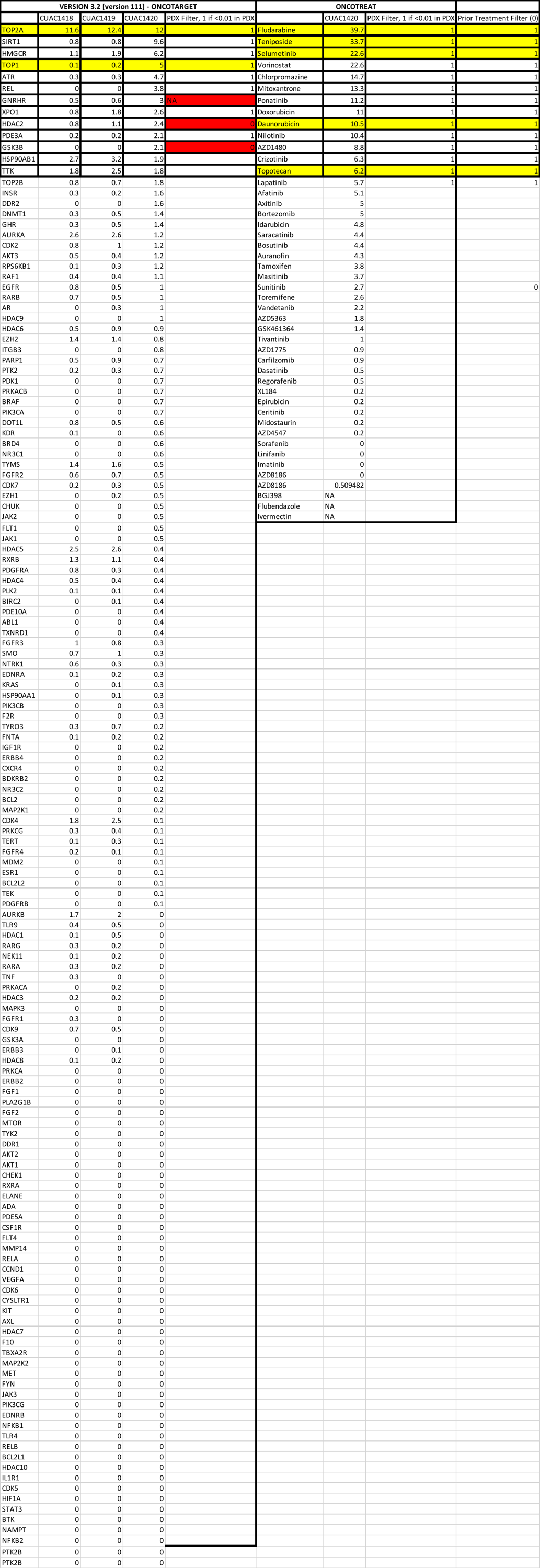

BC.32398

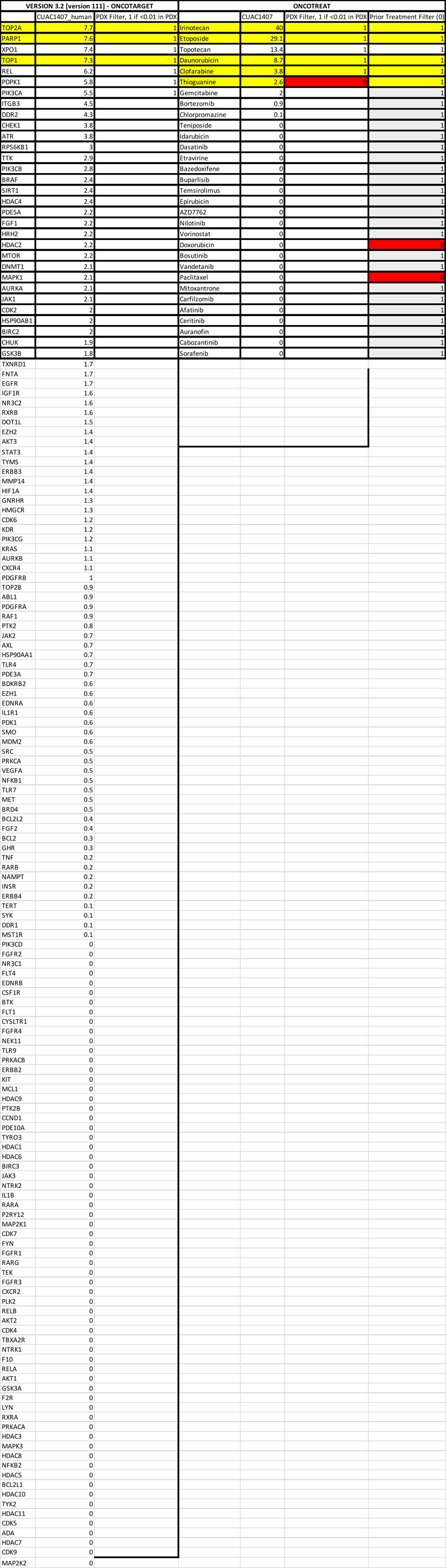

CAR.23659

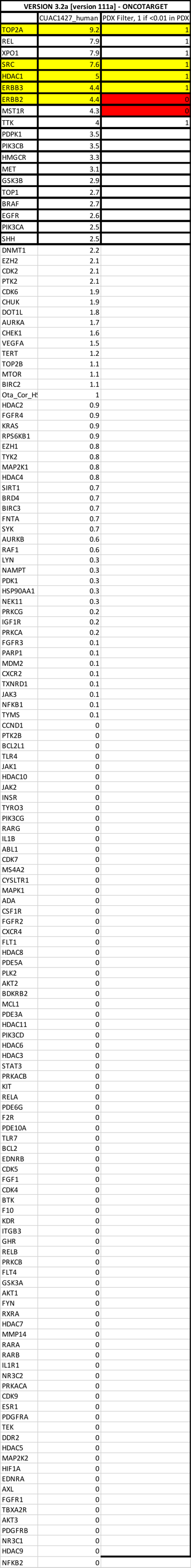

BC.97359

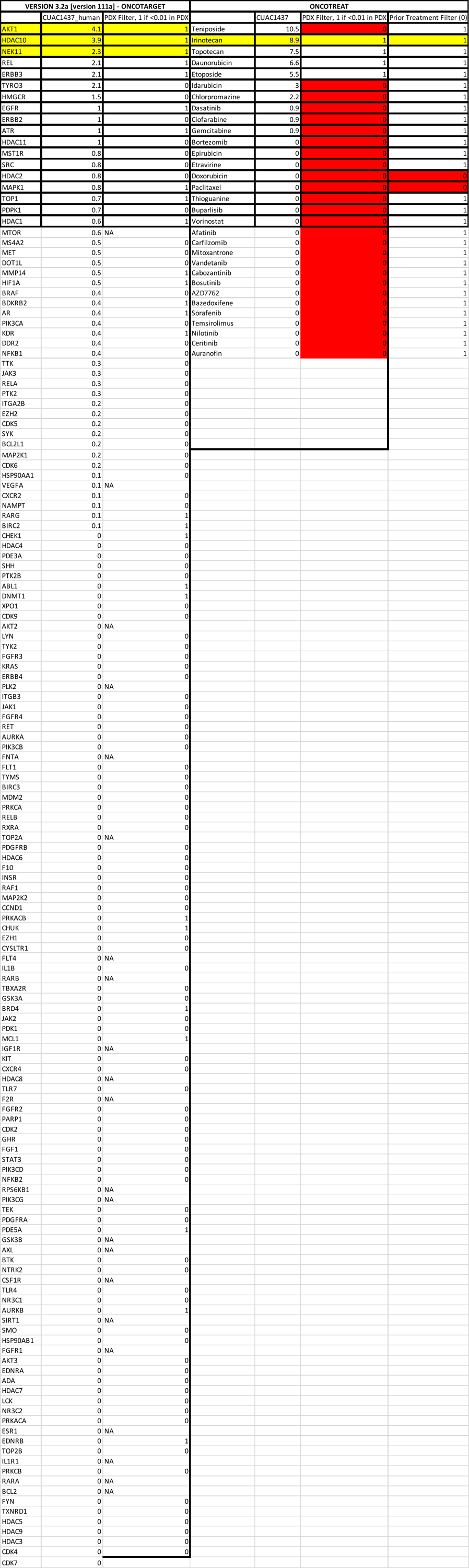

CNS.16474

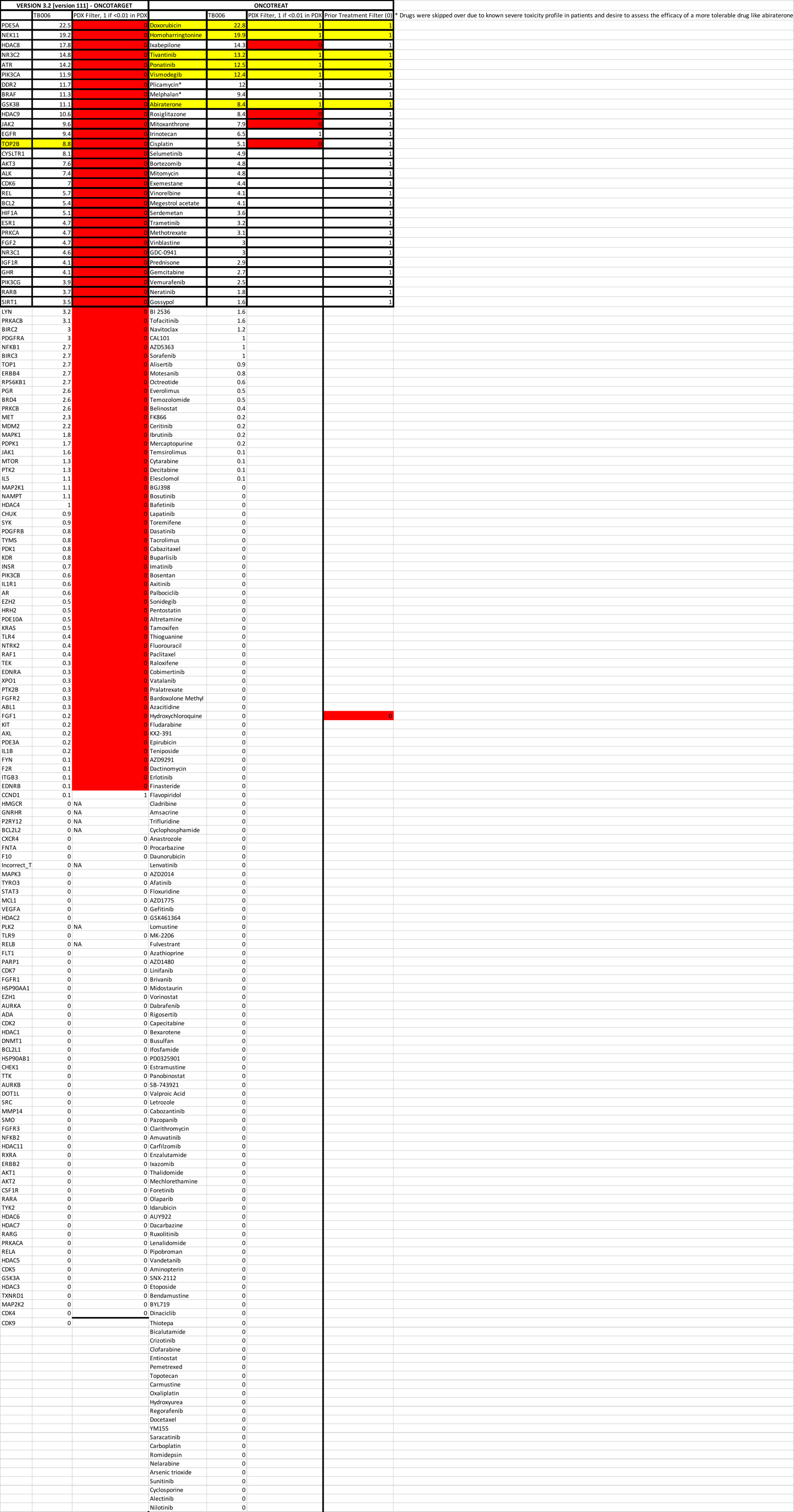

PAC.05647

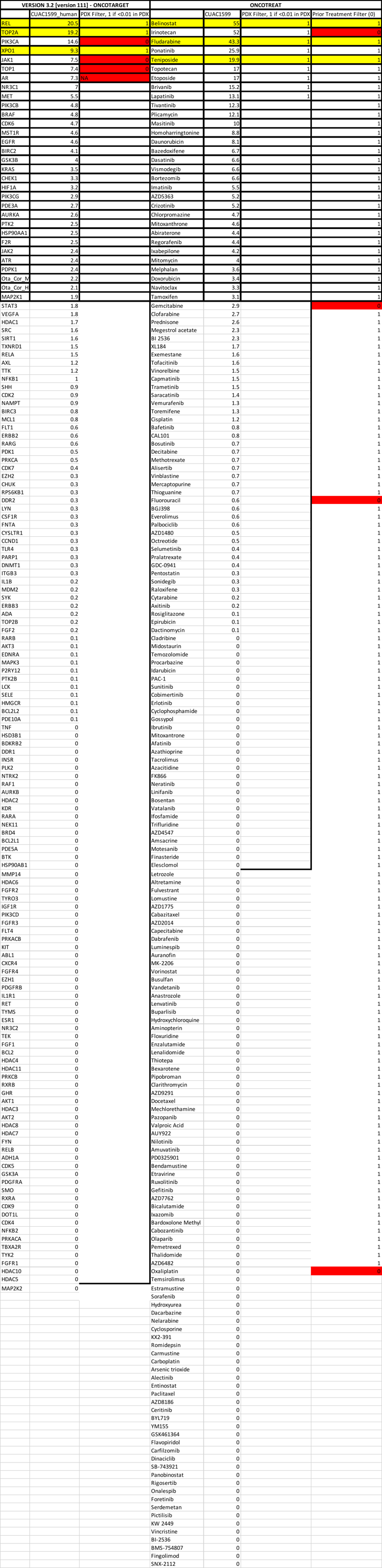

BC.50291

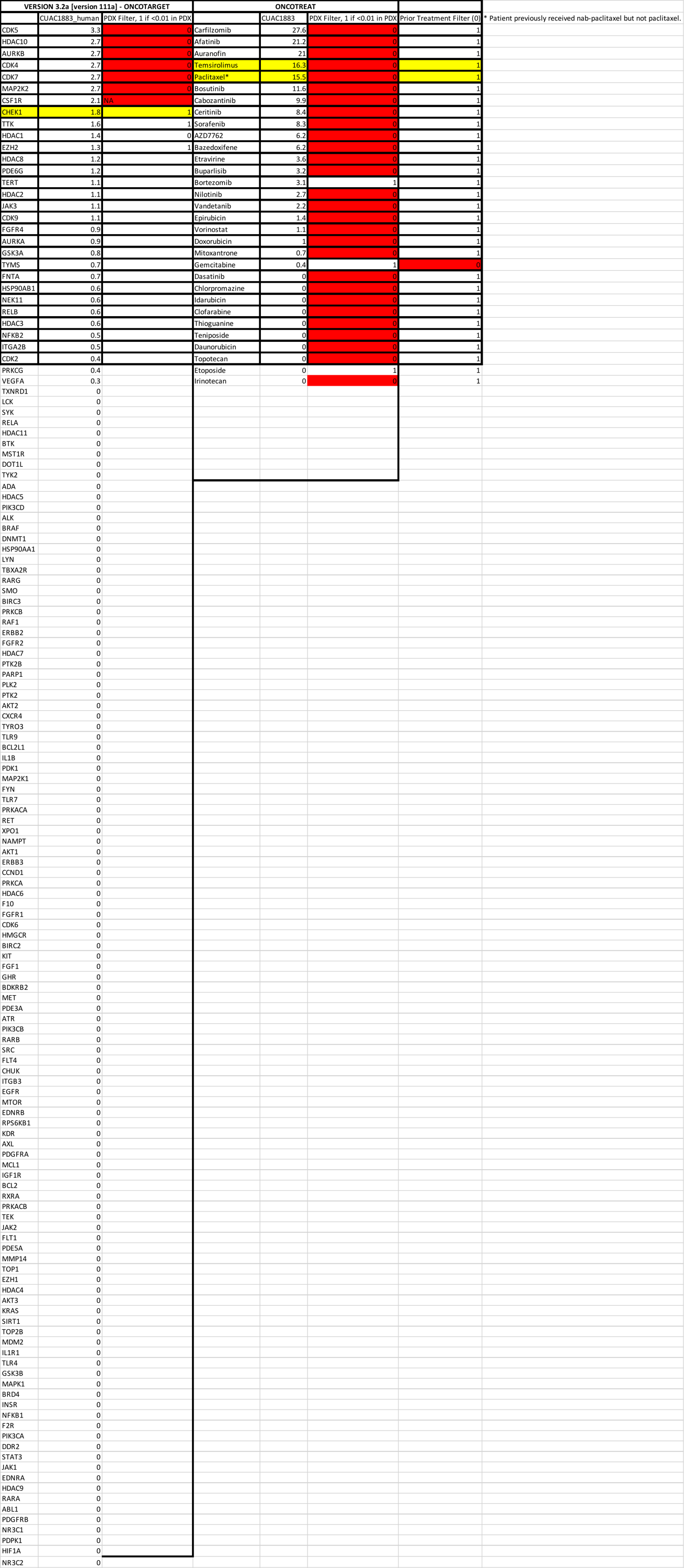

### Table S7

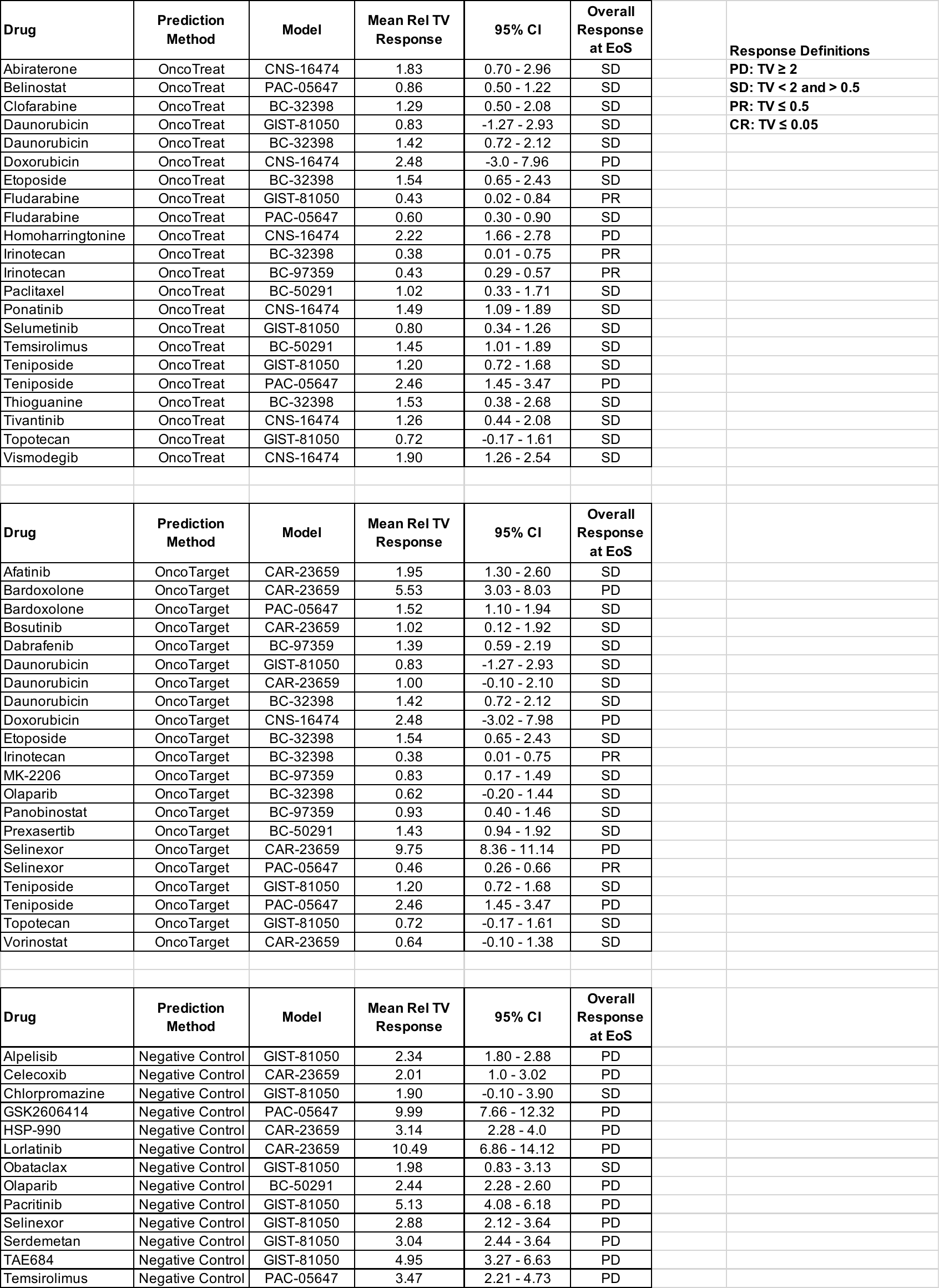
